# Microbial and autoantibody immunogenic repertoires in TIF1γ autoantibody positive dermatomyositis

**DOI:** 10.1101/2020.03.25.007534

**Authors:** Spyridon Megremis, Thomas D. J. Walker, Xiaotong He, James O’Sullivan, William E.R. Ollier, Hector Chinoy, Neil Pendleton, Antony Payton, Lynne Hampson, Ian Hampson, Janine A. Lamb

**Author notes:** These authors contributed equally. These authors contributed equally (Ian Hampson, Janine A. Lamb).

## Abstract

We investigate the accumulated microbial and autoantigen antibody repertoire in adult-onset dermatomyositis patients sero-positive for TIF1γ (TRIM33) autoantibodies. We use an untargeted high-throughput approach which combines immunoglobulin disease-specific epitope-enrichment and identification of microbial and human antigens. Increased microbial diversity was observed in dermatomyositis. Viruses were over-represented and species of the Poxviridae family were significantly enriched. The autoantibodies identified recognised a large portion of the human proteome, including interferon regulated proteins; these proteins were clustered in specific biological processes. Apart from TRIM33, autoantibodies against eleven further TRIM proteins, including TRIM21, were identified. Some of these TRIM proteins shared epitope homology with specific viral species including poxviruses. Our data suggest antibody accumulation in dermatomyositis against an expanded diversity of microbial and human proteins and evidence of non-random targeting of specific signalling pathways. Our findings indicate that molecular mimicry and epitope spreading events may play a significant role in the pathogenesis of dermatomyositis.

## Introduction

The idiopathic inflammatory myopathies (IIM) are a heterogeneous spectrum of rare autoimmune musculoskeletal diseases characterised clinically by muscle weakness and systemic organ involvement. IIM are thought to result from immune activation following environmental exposures in genetically susceptible individuals. Viral and bacterial infections have been reported in individuals with IIM(Miller et al., 2018), but their role in disease pathology is unclear.

Autoantibodies are a key feature of IIM, in common with other autoimmune rheumatic diseases such as rheumatoid arthritis, systemic lupus erythematosus and systemic sclerosis. Myositis-specific autoantibodies, present in approximately 60%–70% of individuals with IIM, are directed against a range of cytoplasmic or nuclear components involved in key intracellular processes, including protein synthesis and chromatin re-modelling(Betteridge et al., 2019). These myositis-specific autoantibodies are often associated with particular clinical features. Individuals with autoantibodies targeting the cytoplasmic nucleic acid sensor MDA5 (also called interferon induced with helicase C domain 1, IFIH1) can present with rapidly progressive interstitial lung disease which is associated with high mortality(Abe et al., 2017; Betteridge et al., 2019). Several members of the tripartite motif (TRIM) protein family are known autoantibody targets in IIM, and there is a strong temporal association between adult-onset dermatomyositis and malignancy onset in individuals with antibodies to transcription intermediary factor 1γ (TIF1γ, TRIM33)(Oldroyd et al., 2019). Many TRIM proteins are important immune regulators(van Tol et al., 2017; Versteeg et al., 2014), and dysregulation of TRIM proteins leading to reduced ability to restrict viral infection has been reported in several autoimmune diseases, including systemic lupus erythematosus and inflammatory bowel disease(Oke et al., 2009; Zhou et al., 2018).

The inducible type I interferon (T1-IFN) cytokine system, part of the innate immune response, also plays a role in autoimmune rheumatic diseases including IIM(Muskardin and Niewold, 2018). The T1-IFN antiviral response is initiated when pathogen-associated molecular patterns are recognized by host pattern recognition receptors and cytosolic receptors for viral nucleic acid. This broad viral recognition process triggers downstream signalling pathways which lead to interferon transcription, protein production and expression of interferon-induced genes; this further enhances the anti-viral machinery(Muskardin and Niewold, 2018). Such a response is critical for host protection against pathogen expansion, and limits infection during the window of time needed to mount an effective specific (adaptive) immune response.

Improved understanding of IIM pathogenesis is required to improve both patient stratification and disease management. In adult-onset dermatomyositis, we propose the following potential molecular mechanism of disease pathogenesis: anti-TIF1 autoantibodies reduce ability to restrict viral infection, which leads to either an increased susceptibility to a wider diversity of viral pathogens or increased exposure to specific anti-TIF1-related viruses. To test this hypothesis, we applied a novel high-resolution and high-throughput comparative screening pipeline (Serum Antibody Repertoire Analysis, SARA) [manuscript in preparation] to both anti-TIF1 autoantibody-positive dermatomyositis patients and matched healthy control plasma. Using this approach, high-throughput antigen epitope-sequencing was integrated with bioinformatic modules to de-convolute accumulated immunogenic responses against the total microbial ‘exposome’ (including viruses, bacteria, archaea and fungi) and human proteins. We report the identification of disease-specific microbial and human protein epitopes which have clinical and aetiological relevance to anti-TIF1 autoantibody-positive dermatomyositis.

## Results

To describe the accumulated antibodies present in dermatomyositis we use the SARA pipeline which integrates an Escherichia coli FliTrx^TM^ random 12 amino acid (AA) peptide display system with epitope signature enrichment through competitive bio-panning and high-throughput DNA sequencing (Figure 1). Competitive bio-panning was applied to pooled total immunoglobulin fractions (IgA, IgG, IgM) purified from the plasma of twenty anti-TIF1 positive adult-onset dermatomyositis patients (DM) and twenty healthy controls (HC) (Table S1). Four sample pools were generated and paired: the first pair contained 10 pooled samples (P10) of DM (DM P10) used for competitive biopanning against 10 pooled samples of HC (HC P10) (Table S1). The second pair included 20 pooled samples (P20) of DM (DM P20) used for competitive biopanning against 20 pooled samples of HC (HC P20) (Table S1). Figure 1 and figure S1 detail achieved metrics while defining the microbial and autoantibody immunogenic repertoires in DM and HC. We retrieved ≈36 million (DM) and ≈24 million (HC) next generation sequencing (NGS) reads which represent 8.7 million (DM) and 4.7 million (HC) expressed epitopes. Cohorts presented highly enriched epitope sequences that are unique to DM (15,522) and HC (4,817) respectively. Our epitope cohorts retrieved 6.75 million (DM) and 2.25 million (HC) microbial or human protein annotations respectively which mapped to 9,111 (DM) and 4,994 (HC) *Distinct* species. 6,202 *Unique* species were identified in DM while 2,085 *Unique* species were identified in HC.

**Figure 1:**
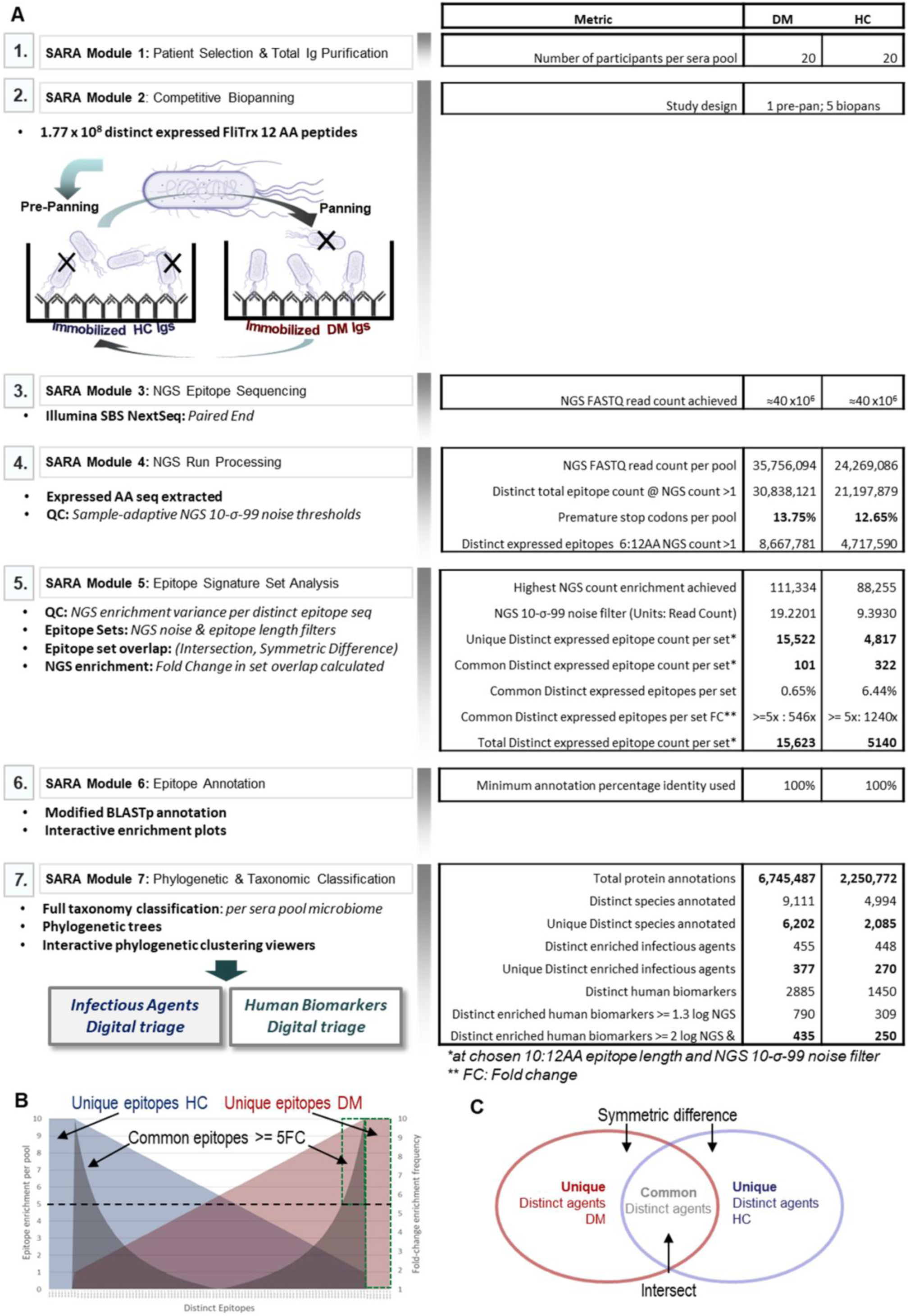
The serum antibody repertoire analysis (SARA) pipeline employed for dermatomyositis patients. (A) The SARA pipeline modules with corresponding technical metrics achieved during the Myositis investigation. Module 1: Patient Selection & Total Ig Purification. Module 2: Competitive Biopanning. Pre-panned clones which bind immobilised healthy control (HC) Igs were discarded whilst unbound clones were immediately passed across immobilised dermatomyositis (DM) pool Igs. Retained clones contain amino acid (AA) sequences representing epitopes of interest. Module 3: Next generation sequencing (NGS) Epitope Sequencing. Enriched epitopes following biopanning were DNA sequenced by a custom Illumina NGS NextSeq process. Module 4: NGS Run Processing. DNA variance regions which code expressed peptide sequences were extracted while NGS noise was controlled by the “10-σ-99” threshold. Module 5: Epitope Signature Set Analysis. Epitope sequence frequency per AA length was determined. Set analysis of distinct epitopes produced unique distinct epitope pools per DM and HC pools respectively (symmetric difference) whilst distinct epitopes common to both cohorts (intersect) were further segregated by a five-fold-change NGS enrichment threshold. Module 6: Epitope Annotation. Our modified BLASTp process was used to annotate epitope sequences to every known protein. Module 7: Phylogenetic & Taxonomic Classification. Annotated Organisms were anchored at appropriate taxonomic classifications to contextualise the plasma pool microbial profiles in the DM and HC cohorts. (B) Example schematic of enriched Distinct epitope sequences (blue: HC pool; Red: DM pool, grey: fold-change). Unique Distinct sequence sets were populated from epitopes found exclusively in one pool or the other (flanking blue or red areas with no fold-change overlay). Common Distinct sequence sets were populated from any epitope that was common to both pools and had a minimum five-fold change enrichment (black horizontal dotted line). Green dotted boxes thus represent the sum of the Unique Distinct and Common Distinct epitope sets that comprised the DM analysis (proportions not to scale). (C) Venn diagram of Unique Distinct (symmetric difference) and Common Distinct (intersect) infectious agents.

### Higher microbial diversity in dermatomyositis

In the DM group, linear epitopes were identified for a total of 1,560 microbial species (Figure S2A) compared to 1,498 in the HC group (Figure S2B). In both groups the highest level of richness (number of different species) was observed for bacteria, followed by viruses, fungi and archaea (Figure S2A & S2B). We defined distinct epitopes as being any discrete epitope sequence retrieved from the DM and HC groups. Unique distinct epitopes represent any discrete epitope sequences which were observed exclusively in either DM or HC. Common distinct epitopes were discrete epitope sequences that were observed in both DM and HC groups by sequence and at a minimum of five-fold the NGS enrichment frequency in one group vs the other group. Unique distinct epitopes represented the vast majority of sequences observed (Fig 1). Based on the distribution of the corresponding microbial species in the pools of 10 and 20 the majority of distinct species are shared between the P20 and P10 samples, whereas the majority of unique species are observed either in P20 or in P10, in both DM and HC (Figure S2C & S2D, respectively). These data demonstrate the presence of a stably-enriched microbial component in DM (Figure S2A) and also in HC (Figure S2B).

The increase in the number of plasma samples within the pool used for cross-panning, from a pool of 10 (P10) to a pool of 20 (P20), had a differential effect between the two groups: in DM, an increase in the number of microbial species was observed from 728 (DM P10) to 832 (DM P20), whereas in the HC group a decrease was observed from 780 (HC P10) to 718 (HC P20) species. The number of epitopes per microbial species in both the DM and the HC significantly increased with increasing pooling size (Figure 2A). The number of NGS reads per microbial species significantly increased in the DM P20 (95% CI: 3.97-4.13) compared to DM P10 (95% CI: 3.90-3.98), whereas it did not change in the HC P20 (95% CI: 4.09-4.26) compared to the P10 (95% CI: 4.34-4.43) (Figure 2B).

**Figure 2:**
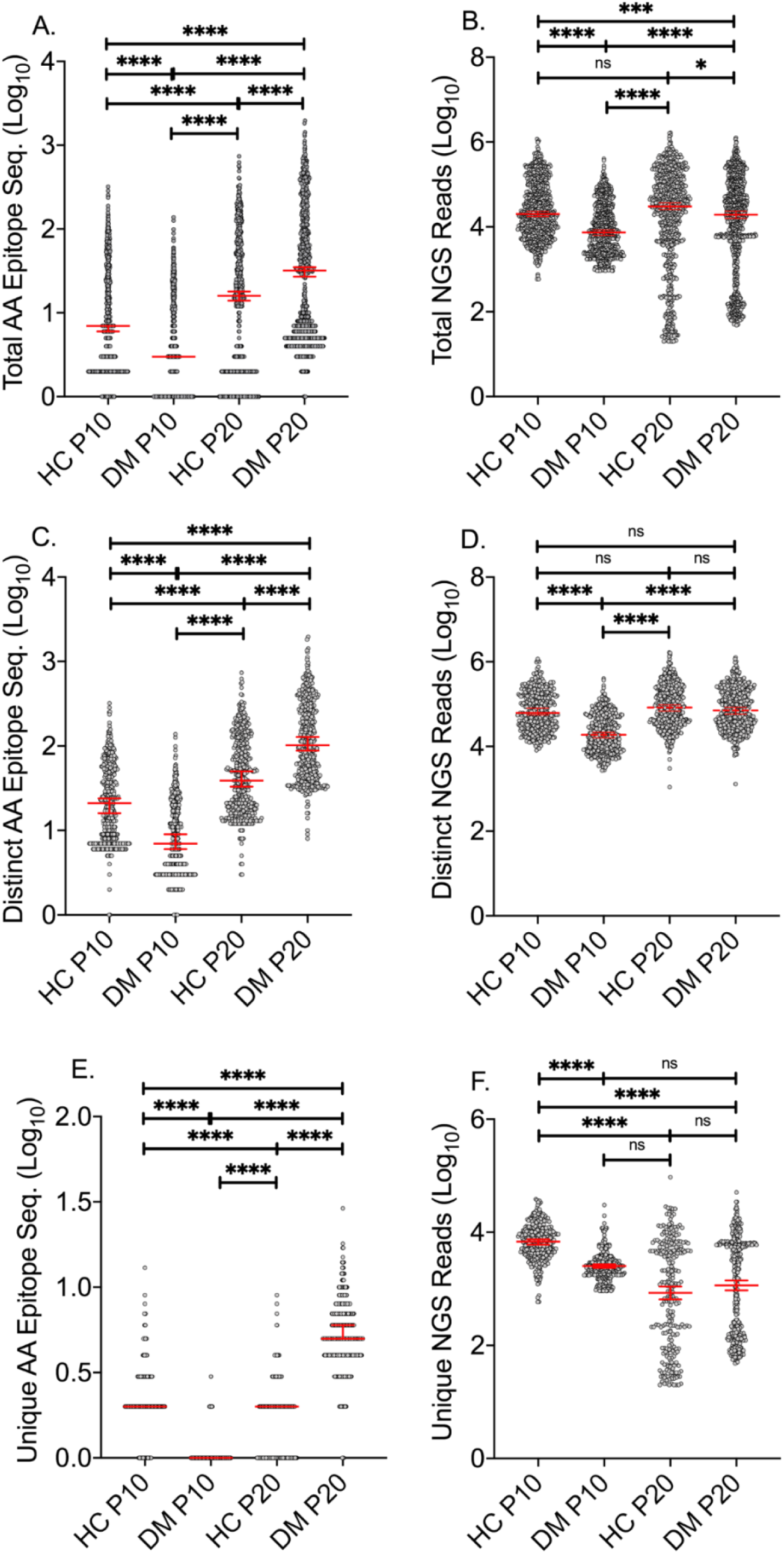
Microbial amino acid epitopes and their abundance in dermatomyositis and healthy controls. The number of different epitopes per microbial species (left panel) and the number of next generation sequencing (NGS) reads that mapped to these epitopes (right panel) are depicted; (A) Total number of microbial amino acid (AA) epitopes, (B) Microbial NGS reads, (C) Number of distinct AA epitopes, (D) Distinct microbial NGS reads, (E) Number of unique AA epitopes, and (F) Unique microbial NGS reads. All values were log_10_ transformed. Dunn’s multiple comparisons test was used to test specific sample pairs. Differences were significant at the 0.05 level; * for p<0.05, ** for p<0.01, *** for p<0.001 etc. DM, dermatomyositis; HC, healthy controls; ns: not-significant; P10: pool of 10; P20: pool of 20. The median with 95%CI is annotated.

The number of distinct epitopes per microbe (epitopes present in both groups at minimum five-fold enrichment) significantly increased in the P20s compared to P10s in both groups (Figure 2C); (n=432, 95%CI: 0.86-0.94) to (n=455, 95%CI: 2.02-2.10) in DM and 432 (n=432, 95%CI: 1.29-1.37) to 448 (n=448, 95%CI: 1.60-1.68) in HC. The unique microbial epitopes which were identified only in one of the two groups significantly increased in DM P20 (n=377, 95%CI: 0.69-0.77) compared to DM P10 (n=296, 95%CI: 0.01-0.03) and decreased in HC P20 (n=270, 95%CI: 0.18-0.23) compared to HC P10 (n=348, 95%CI: 0.31-0.36) (Figure 2E). Collectively, these data indicate an expansion of the DM-specific microbial diversity with increasing DM plasma sample size, whereas the opposite effect was observed in HC.

The number of enriched NGS reads per distinct microbial species significantly increased in DM P20 (95%CI: 4.83-4.92) compared to DM P10 (95%CI: 4.27-4.36), while no significant difference was observed in HC P20 (95%CI: 4.89-4.98) compared to HC P10 (95%CI: 4.80-4.89) (Figure 2D). The number of sequencing reads per unique microbial species did not differ significantly between DM P20 and DM P10 (adjusted p value >0.999), however it significantly decreased in the HC P20 (95%CI: 2.81-3.04) compared to the HC P10 (95%CI: 3.78-3.85) (Figure 2F). Overall, the differential effect of increased sample size on the observed microbial exposure between DM and HC suggests there is a higher biological variability in DM than in HC.

DM P20 exhibited a significantly higher number of microbial AA epitopes per microbial species compared to the HC P20 (Figure 2A) and a higher number of total identified microbial species (832 versus 718). The same difference was observed in both distinct (Figure 2C) and unique epitope sequences (Figure 2E). Overall, we demonstrate that plasma from anti-TIF1 DM patients contain a higher number of microbial epitopes per species and against a wider microbial repertoire.

### Significant role of viruses in dermatomyositis

We focused our further analysis on the DM P20 and HC P20 subgroups. To stratify the microbial species based on their potential significance in DM, we first normalised the number of NGS reads against the total number of epitopes per microbial species (NGSR_e-norm_) (Figure S3A-S3D), and studied the relative ranking of viruses and cellular microbes (Figure 3A & 3B). Secondly, we evaluated the mean NGSR_e-norm_ at the viral family taxonomy level (Figure 4D & 4E), and, thirdly, we recorded the total number of species contributing to each family (Figure 4D & 4E) (Table S2). Thus, we tested for potential high taxonomy-level organisation of microbes in DM. Ranking the microbial species based on decreasing NSGR_e-norm_ we observed that viruses were over-represented in the top 10% of dominant microbial species relative to the total number of viral species present in both DM and HC (Figure 3A & 3B). Specifically, 18.47% and 17.93% of viral species were present in the top 10% of microbes in the DM P20 and HC P20, respectively (Figure 3C & 3D). Cellular microbes were under-represented in the dominant species with 5.95% in DM P20 and 7.58% in HC P20 (Figure 3C & 3D). The over-representation of viral compared to cellular microbial species in the top-ranked microbes enriched in DM and HC suggests an important role of virus exposure in the environment-host immune cross-talk in both healthy and disease states, and implies that viruses may play an important role in the pathology of DM.

**Figure 3:**
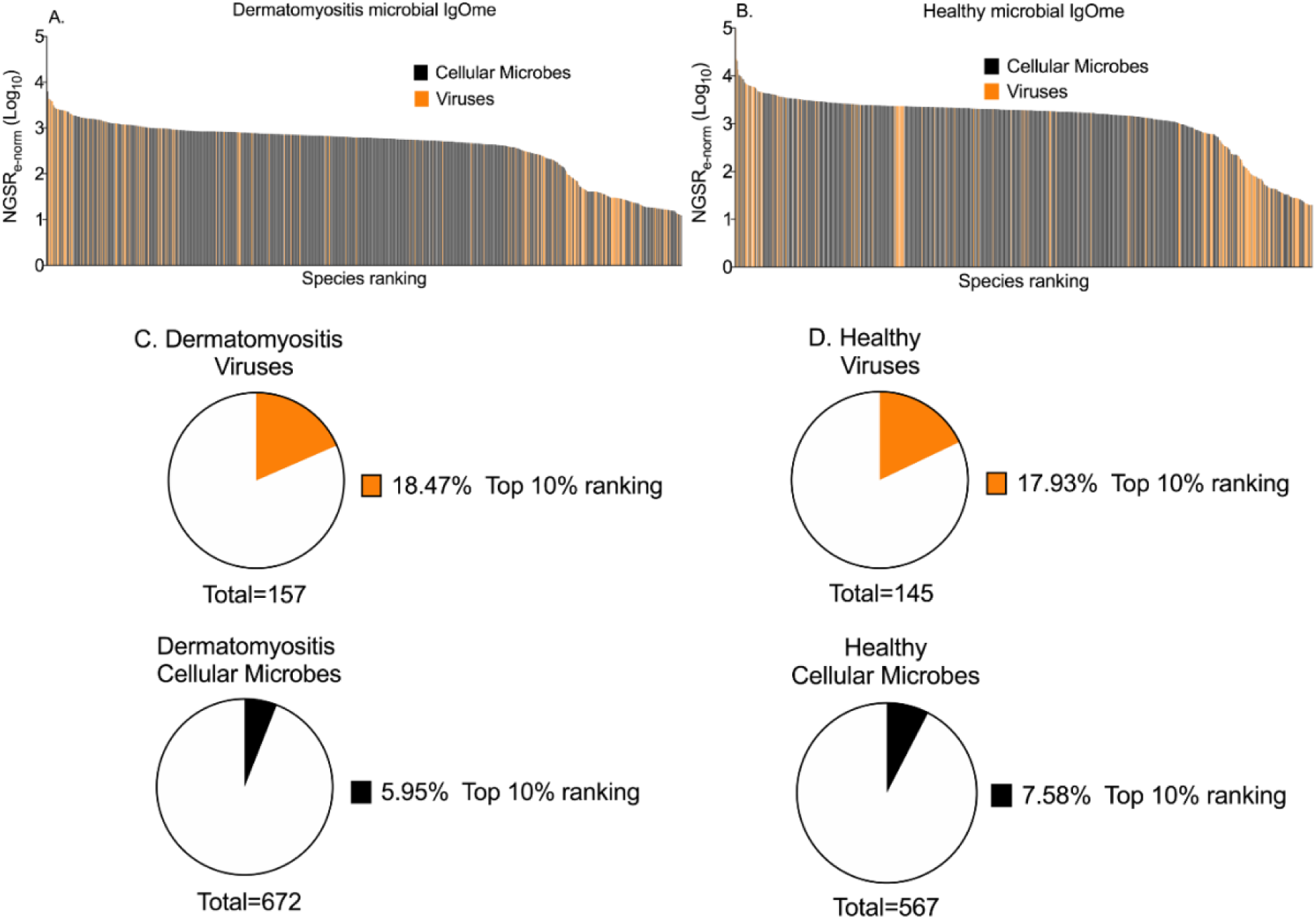
Ranking of identified microbial species based on the number of NGS reads and number of epitopes. Microbial species were grouped in cellular microbes (Bacteria, Eukaryota, and Archaea) and viruses and ranked based on the NGSR_e-norm_. The distribution of members of the two groups can be seen in (A) dermatomyositis, and (B) healthy controls; Viruses were superimposed over cellular microbes. The pie charts depict the proportion of each microbial group that is represented in the top 10% of ranked species in (C) dermatomyositis and (D) healthy controls. Orange: viruses, Black: cellular microbes.

**Figure 4:**
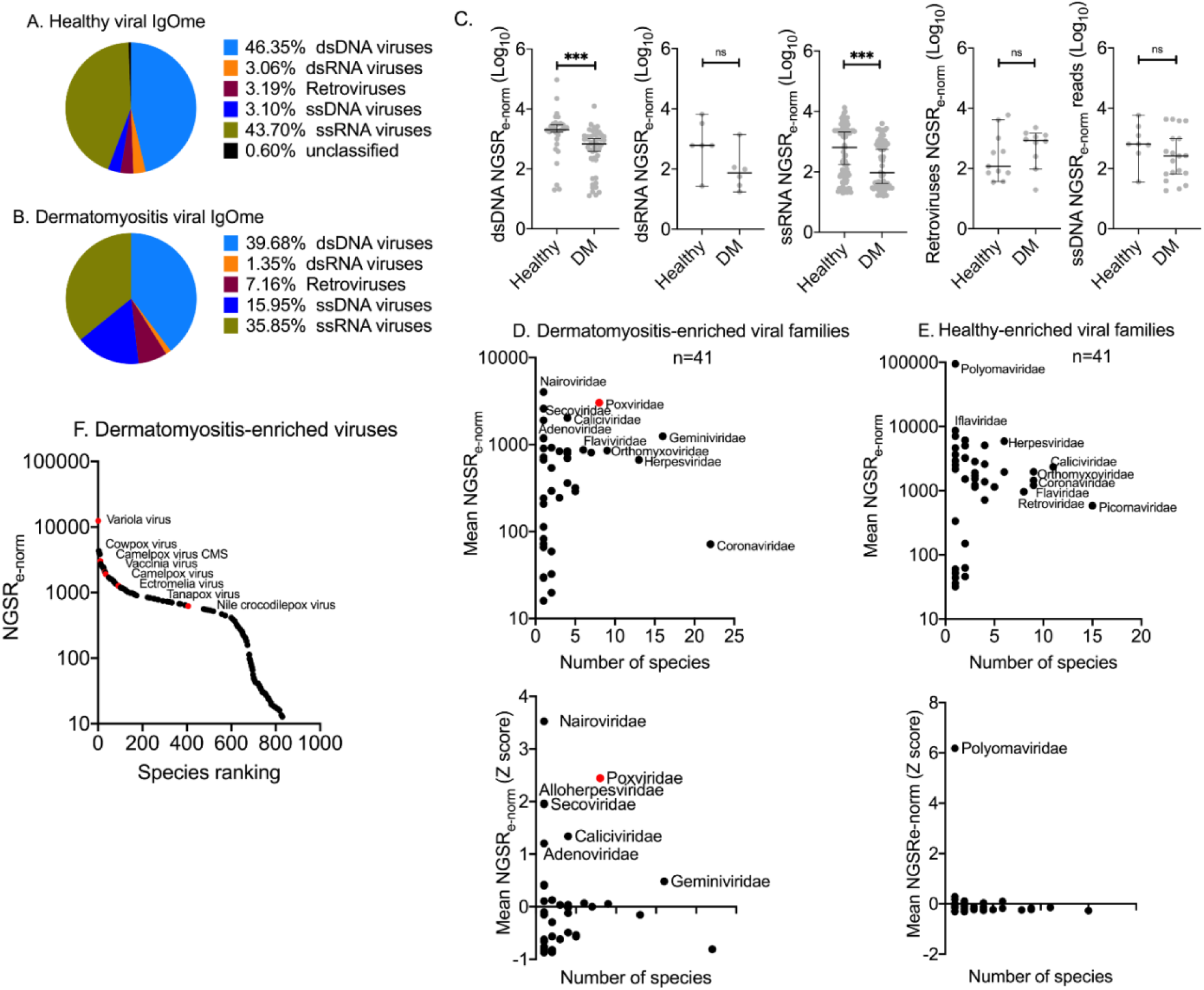
The dermatomyositis-specific viral IgOme. Viral species were grouped based on genome type into 6 groups. The contribution of each viral group to the total NGSR_e-norm_ output was evaluated in (A) healthy control (HC) P20, and (B) dermatomyositis (DM) P20. (C) Scatter plots of viral normalised abundance, mean NGSRe-norm, in DM and HC. Viruses are grouped based on their genome type. Scatter plots of viral family richness (number of species per family) and mean NGSRe-norm in: (D) DM, and (E) HC; In the upper panel NGSR_e-norm_ values are depicted. In the lower panel the NGSR_e-norm_ values are Z transformed to identify viral targets with high NGSR_e-norm_ (outliers). The Poxviridae viral family is depicted with red colour. (F) Scatter plot of species ranking based on NGSRe-norm in DM; Species of the Poxviridae family are depicted with red colour. Differences were significant at the 0.05 level; * for p<0.05, ** for p<0.01, *** for p< 0.001. The median with 95%CI is annotated.

### Poxviruses are tightly linked with dermatomyositis

Viruses were grouped in high-order taxonomic groups based on their genome type (Figure 4A & 4B). The HC viral IgOme profile contained a higher proportion of total double stranded DNA (dsDNA) (NGSR_e-norm_ 46.35% vs. 39.68%) and single stranded RNA (ssRNA) (NGSR_e-norm_ 43.70% vs. 35.85%) than DM (Figure 4A & 4B). The DM viral IgOme contained a higher proportion of ssDNA viruses (NGSR_e-norm_ 15.95% vs. 3.10%) and RNA reverse-transcribing retroviruses (NGSR_e-norm_ 7.16% vs. 3.19%) (Figure 4A & 4B). To test whether these observations were due to a generalised differentiating signal amongst viral species within each viral group we compared the species-specific NGSR_e-norm_ per viral category between DM and HC (Figure 4C). The dsDNA and ssRNA NGSR_e-norm_ were elevated in the HC plasma compared to DM, whereas no significant change was observed in ssDNA viruses or RNA reverse-transcribing viruses (Figure 4C).

Overall, 47.9% (23 out of 48) of viral families were represented in both groups. We compared the viral IgOme at the family level, and we recorded the mean NGSR_e-norm_ with increasing enrichment for each viral family (Figure 4D & 4E). In DM, the five richest viral families were Coronaviridae (20 species), Geminiviridae (16 species), Herpesviridae (13 species), Orthomyxoviridae (9 species) and Poxviridae (8 species) (Figure 4D) (Table S2). Of these, Geminiviridae infect plants, whereas all other viruses can infect and cause disease in humans. In the HC, Picornaviridae (15 species) was the most enriched virus group followed by Caliciviridae (11 species), Orthomyxoviridae (9 species), Coronaviridae (9 species), and Retroviridae (8 species) (Figure 4E) (Table S2).

We evaluated the mean NGSR_e-norm_ for each family. In DM, Nairoviridae (n=1), Poxviridae (n=8), Secoviridae (natural host: plants) (n=1), Caliciviridae (n=4) and Adenoviridae (n=1) were the top 5 ranked families (Table S3). Poxviridae was the one family that ranked highly regarding both the NGSR_e-norm_ (Table S3) and the family richness (n=8, 95%CI: 2.98-3.61) (Table S2). All 8 pox viruses had high NSGR_e-norm_ and were amongst the dominant DM viral species (Figure 4F). Specifically, Variola virus had the highest NSGR_e-norm_ amongst all identified viral species (Figure 4F). In the HC P20, Polyomaviridae (n=1), Iflaviridae (n=1), Podoviridae (n=1), Tymoviridae (n=2), and Myoviridae (n=3) were the viral families with the highest mean NGSR_e-norm_. Podoviridae and Myoviridae are prokaryotic viruses infecting bacteria, Tymoviridae infect plants and Iflaviridae infect insects (ViralZone root-ExPASy)(Hulo et al., 2011). We also used Z transformation of NGSR_e-norm_ to define the precise location and rank of each viral family within the DM and HC distribution (Figure 4D & 4E). In DM, multiple viral families (Nairoviridae, Poxviridae, Secoviridae, Alloherpesviridae, Caliciviridae and Adenoviridae) with high NGSR_e-norm_ Z score occurred with at least one standard deviation above the group mean. In HC, only Polyomaviridae (n=1) occurred with high NGSR_e-norm_ Z score. In Figure S4 we provide a cladogram of DM-specific viral species.

### Detection of multiple TRIM proteins in dermatomyositis

We queried the NGS reads against the human proteome to identify accumulated autoantibodies in the pooled plasma. In DM P20 we identified TRIM33 (Mean log_10_: 1.301), in accordance with the autoimmune profile of our selected plasma (Figure 5A) which was not present in HC P20. We also identified 11 additional TRIM proteins in DM P20 which were TRIMs 21, 69, 47, 46, 27, 60, 10, 7, 77, 3 and TRIML2 (Figure 5A). Of these, TRIM21 was also detected in the healthy samples but with significantly lower NGS reads, and TRIM25 (Mean log_10_: 1.778) was observed only in the HC P20 (Figure 5A). These results confirm the presence of TRIM33 autoantibodies in the selected twenty anti-TRIM33-positive DM patients and demonstrate the presence of autoantibodies against other members of the TRIM protein family exclusively in DM.

**Figure 5:**
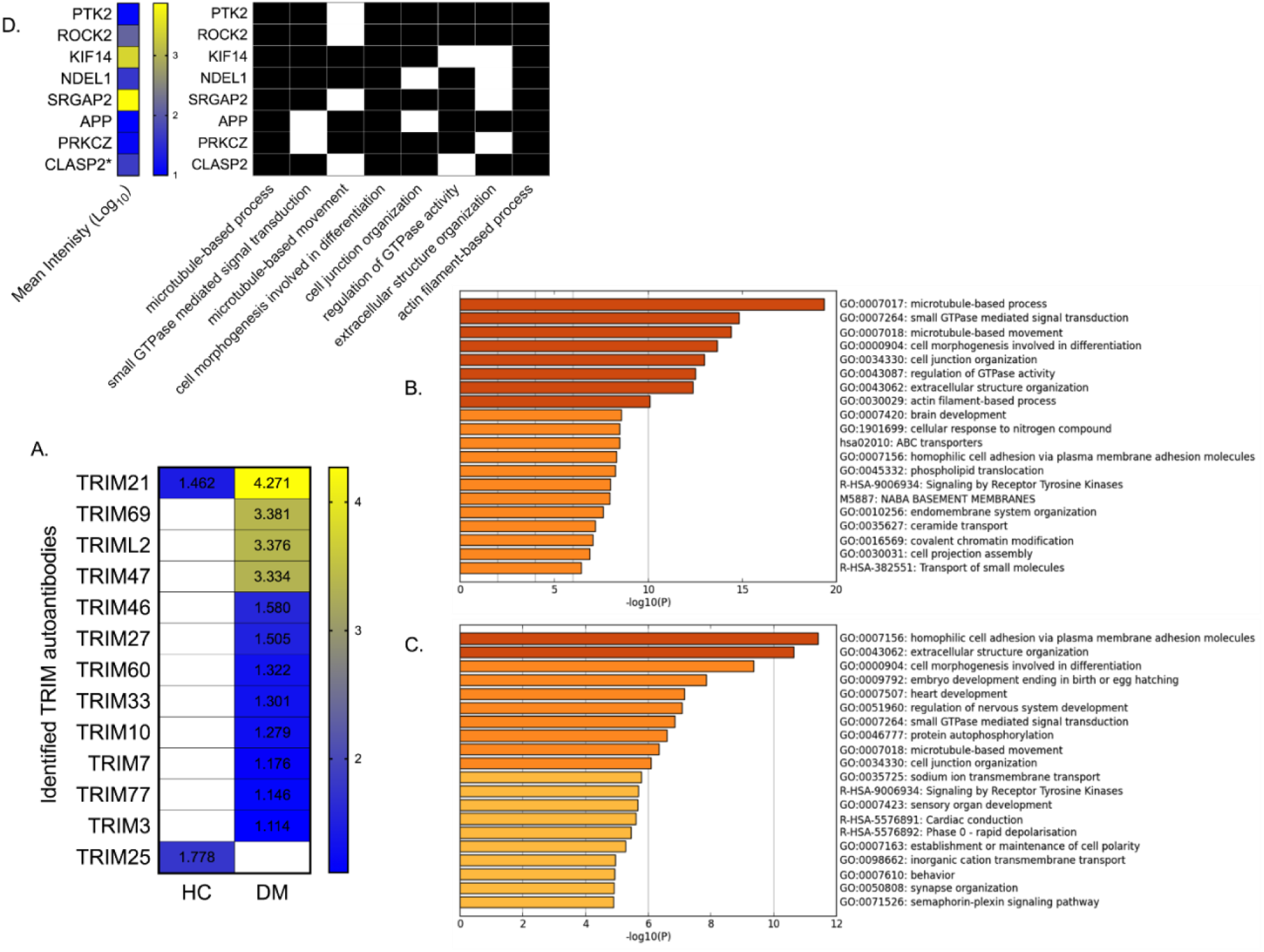
TRIM autoantibodies identified in Dermatomyositis. (A) Heatmap of mean log_10_ intensities of identified TRIM proteins in healthy control (HC) and dermatomyositis (DM). (B) Gene ontology processes significantly enriched in (B) DM, and (C) HC. (D) Binary colour-coding of presence/absence of the autoantibody protein targets that were enriched in at least 6 out of 8 DM-specific GO processes (presence: black, absence: white). The mean log_10_ intensity of each protein is annotated in a colour coded heatmap (low: blue to high: yellow).

### An expanded dermatomyositis-specific and IFN-regulated human proteome

From our antibody analysis, in DM P20 we identified a total of 2885 human protein targets, of which 2537 were highly specific sequence annotation hits (Mean Sig. < 0.578). In HC P20 we detected 1450 human protein targets, 811 with high specificity (Mean Sig. < 0.55). Autoantibodies identified in both DM and HC constituted only 13% (n=498) of total proteins and 8.4% (n=260) of highly specific proteins suggesting a strong disease-specific proteome signature. The 2:1 (total proteins) and 3:1 (high specificity) ratios of identified autoantibodies in DM over HC, suggest that an expanded subset of the human proteome is targeted by autoantibodies in DM patients compared to HC.

Due to the role of TRIM proteins in IFN signalling, we asked whether the autoantibody protein-targets are regulated by interferons. In the DM dataset 1,560 proteins were predicted to be regulated by IFNs compared to 518 in HC (Interferome v2.01)(Rusinova et al., 2013). The vast majority of these proteins are regulated by interferons type I, type II or both (Figure S5A). Regardless of the IFN type we observed a higher number of IFN-regulated proteins in DM compared to HC suggesting a DM-specific enrichment of immunoglobulins against IFN-regulated human proteins. We focused our search directly for IFN autoantibodies expressed in our samples. In DM, IFNGR1 autoantibodies were present, whereas they were absent in HC (Figure S5B). We expanded our search for autoantibodies against known proteins that are highly ranked within the IFNG signalling pathway (gene rank within the SuperPath; GeneCardsSuite: PathCards)(Belinky et al., 2015). Autoantibodies against 26 IFNG-related proteins were observed in DM, whereas only 5 were found in HC (Figure S5B). The protein-protein interaction (PPI) enrichment score in DM was <1.0e-16 versus 0.123 in the healthy sample (STRING 11.0)(Szklarczyk et al., 2019) (Figure S5C & 5D). Overall, these data suggest the accumulation of antibodies in DM against proteins that strongly contribute to IFNG signalling and the broader antiviral mechanism by IFN-regulated proteins.

To describe the biological functions of all the identified proteins the Gene Ontology (GO) framework was used (Figure 5B & 5C). In DM P20 eight GO biological processes were highly enriched (Figure 5B). These processes were represented by an average of 25.6% of GO-specific proteins (n=996, 95%CI: 21.2%-29.7%) (Figure S6A). In HC P20, ten GO processes were enriched with an overall lower average coverage of 14.5% (n=592, range: 12.5%-15.7%), as less proteins mapped to the same GO unit. In the top-ranked GO processes enriched in DM, the GO coverage was higher in DM than HC regardless of the GO process (Figure S6A). More than one third (39.2%: 996 out of 2537) of the DM autoantibody targets were part of the 8 identified biological processes; 18.6% of them (186 out of 996) were also present in HC P20. DM processes involved structural elements including microtubule-based processes, actin-filament functions, cell junction organisation, extracellular structure organisation, cell morphogenesis in differentiation and small GTPase mediated signal transduction (Figure 5B). PTK2 and ROCK2 were shared between 7 out of 8 functions, and, KIF14, NDEL1, SRGAP2, APP, PRKCZ, and CLASP2 were shared amongst 6 out of 8 functions (Figure 5D). None of these proteins were observed in HC. The fact that the identified DM-specific and HC-specific proteins were robustly clustered in biological functions suggests non-random targeting of specific signalling pathways by autoantibodies. In DM, these processes share multiple autoantibody protein targets suggesting the presence of a DM-specific autoantibody-targeted proteome module (Figure S6B).

### Shared epitope sequences between Variola virus, Poxviridae, HIV and TRIM proteins

We identified that epitope sequences against the Poxviridae family of viruses were significantly enriched in DM, including variola virus which had the highest NGSR_e-norm_ (Fig. 4F). Given that our DM patients were TRIM33 autoantibody positive and due to the fact that TRIM proteins, IFNG and the IFN antiviral mechanism were significantly enriched in our proteome data, we searched for potential links between Variola virus, other members of the Poxviridae family and TRIM proteins. Since molecular mimicry is a potential mechanism of autoantibody generation, we aligned the identified variola virus and TRIM epitopes; the sequence annotation thresholds that we used in our bioinformatics pipeline guaranteed robust annotation of the epitope sequences. This would affect cross-kingdom epitope alignment, since we have maximized the phylogenetic distance between each microbial epitope and human proteins. To account for this, we started our epitope sequence analysis with the widest possible diversity of identified variola and TRIM epitopes, sacrificing specificity i.e. epitopes with more than 80% match to variola virus, and TRIM epitopes with an identity of more than 50%. The phylogenetic distances are shown in the circular cladogram in Figure 6. We identified two different clades containing leaf nodes of TRIM and variola epitopes of high similarity (branch lengths <0.1). The first clade involved the TRIM3 epitope “RIPDDVRRRPGC” and three additional epitopes “RI(Q)DDVRRRPGC”, “RI(Q)DDV(H)RRPGC” and “RI(Q)DD(V)(S)RRPGC” each of which mapped to VARV GER58 hdlg 202 and VARV GUI69 005 202. The second clade contained the TRIM3 epitope “SSHARYKSVRFS”, and “SSHARYKSVRFS”, “SSHARYKS(M)RFS “SSHARYKSLRFS”, “SSHARYKS(L)RF”, and “SSHARYKSLRF(T)” of a variola virus (unnamed protein product and viral DNA polymerase processivity factor). We reinstated the annotation specificity thresholds for TRIM and variola epitopes to the default levels (strict, high specificity, maximisation of phylogenetic distances) (Figure S7A). The epitope sequence “RI(P)DDVRRRPGC” was retained and shared between TRIM3 (branch length: 0.0541) and VARV GER58 hdlg 202 (branch length: 0.0344). The epitope sequences “SSHARYKSLRFS” and “SSHARYKSLRF” were specific for variola virus but were no longer annotated as TRIM epitopes (Figure S7A). We asked whether the above epitope sequences are shared amongst different Poxviridae species that we identified in DM P20 (Figure 4F). The 3 epitopes (“SSHARYKSLRFS”, “SSHARYKSLRF”, and “RIQDDVRRRPGC”) were shared with high homology between variola, vaccinia, ectromelia, cowpox and camelpox viruses (Figure S7B). We aligned the DM TRIM proteins that we identified to test the conservation level of the above epitopes in TRIMs (Figure 7A). TRIM3 and TRIM33 shared high similarity of AA sequence and seem to diverge from the rest of TRIMs across the region of interest (Figure 7A).

**Figure 6:**
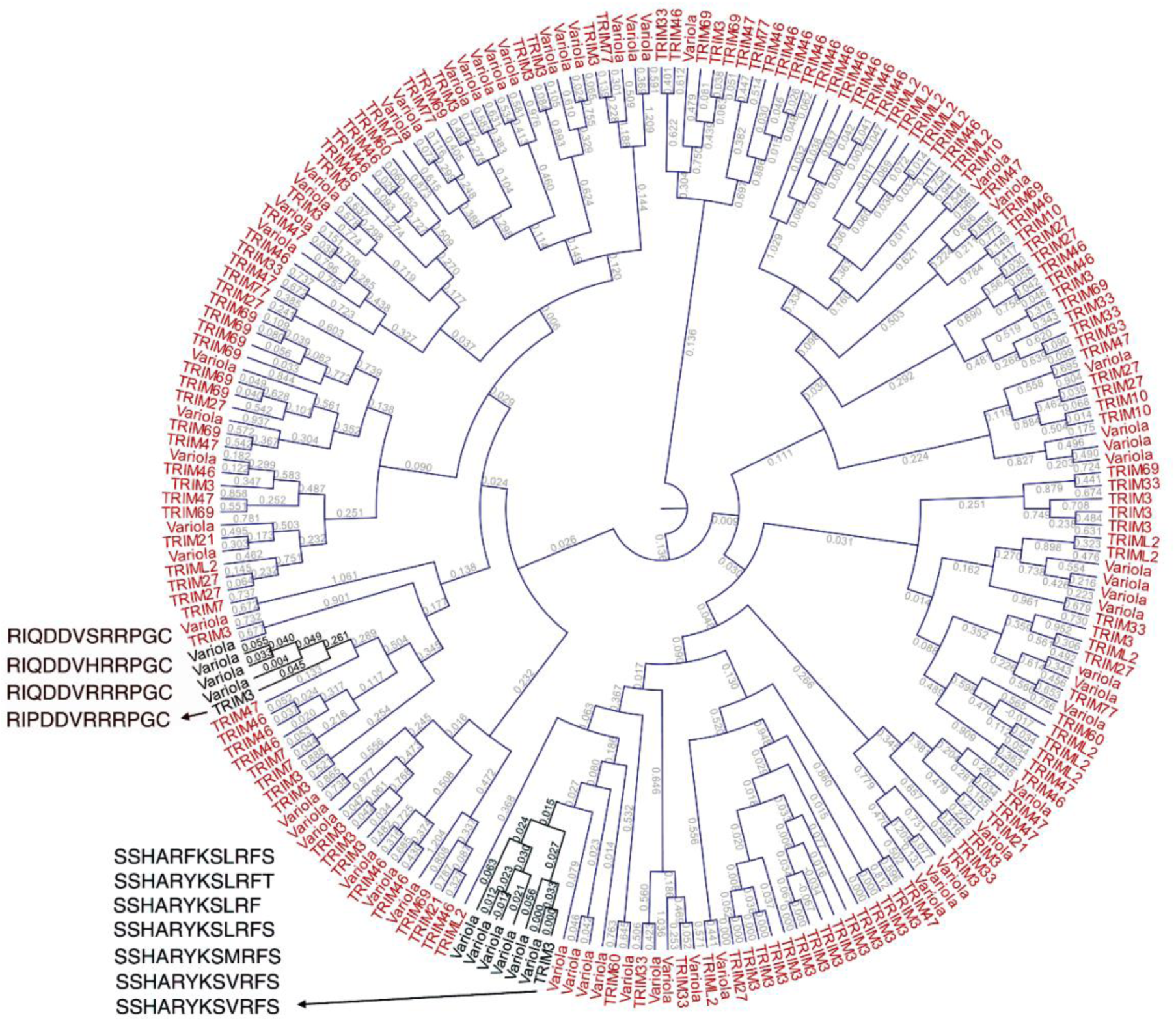
Circular cladogram of phylogenetic distances between variola virus and TRIM epitopes identified in dermatomyositis. The DM-specific epitope sequences identified in TRIM3 and variola virus had the highest similarity and are annotated next to the cladogram clades. Branch lengths represent phylogenetic distance (Kimura protein distance).

**Figure 7:**
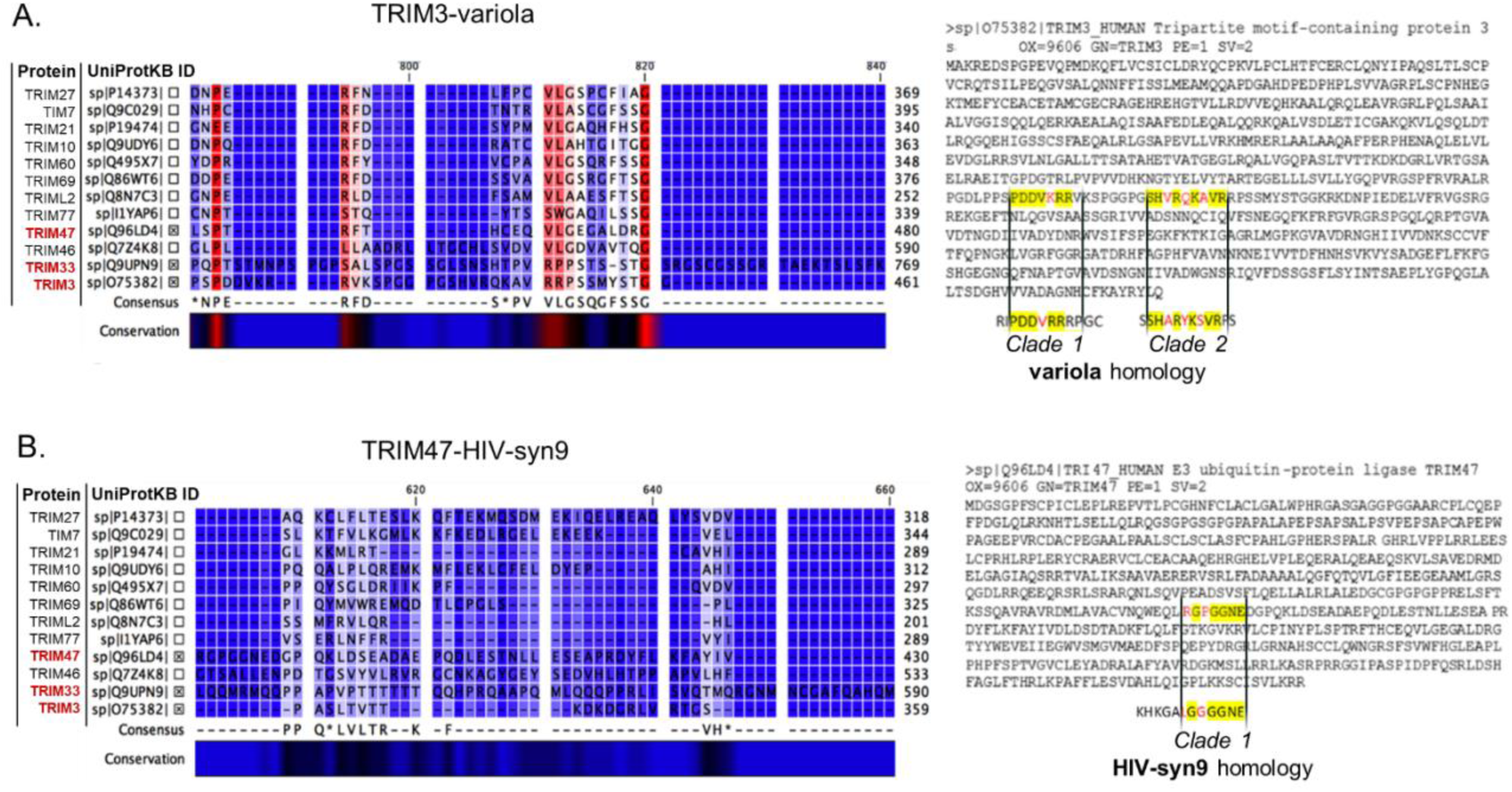
Partial amino acid alignments of the DM-specific TRIM proteins that share amino acid similarity with viral species. Alignment of all identified TRIM proteins in DM (Fig. 5A); TRIM3 is located on the bottom of the alignments plots. Above TRIM3 (sp|O75382) are TRIM33 (sp|Q9UPN9) and TRIM47 (sp|Q96LD4); cross-marked. Conservation level is presented as a colour gradient (high; red, low; blue). (A) Partial alignment of the region where the TRIM3-variola high similarity epitopes were identified. The TRIM3 protein sequence is presented next to the alignment plot. Yellow: the two TRIM3 epitopes with high amino acid (AA) similarity to variola and poxviruses (Figure 6). Black font: identical AAs between TRIM3 and variola virus. Red font; different AAs between TRIM3 and variola virus. Both epitopes are located only 6 AAs apart (relative to TRIM3 protein sequence). TRIM3 presents high similarity with TRIM33 in respect to the specific AA epitopes. (B) Partial alignment of the region where the TRIM47-HIV-syn9 high similarity epitope was identified. The TRIM47 protein sequence is presented next to the alignment plot. Yellow: the TRIM47 epitopes with high AA similarity to HIV and syn9 phage (Figure S7D). Black font: identical AAs amongst TRIM47 and the two viruses. Red font; different AAs amongst TRIM47 and the two viruses. The epitope is poorly conserved amongst the DM-specific TRIM proteins.

Finally, we aligned all DM-enriched viral and TRIM epitopes (Figure S7C) and identified a third epitope “KHKGALGGGGNE” of TRIM47 which is shared by Human immunodeficiency virus 1 “KHKGALGGGG(N)E”, and “KHKG(D)LGGGG(Y)E” shared by Synechococcus phage syn9 “KHKG(D)LGGGG(Y)E” (Figure S7D). The epitope was poorly conserved amongst the DM-specific TRIM proteins (Figure 7B). Overall, we have identified two epitope sequences that are shared between Variola virus and TRIM3 but also between members of the Poxviridae family. A third epitope that is shared between TRIM47, HIV-1 and Syn9 was also observed. These findings suggest that molecular mimicry events, at least amongst these specific viral and TRIM epitopes, is a potential mechanism of pathogenesis in DM.

## Discussion

This is the first study to investigate the accumulated microbial and autoantigen antibody repertoire in adult-onset dermatomyositis (DM) patients positive for antibodies against TIF1 (TRIM33). We used an un-targeted high-throughput approach which combines immunoglobulin disease-specific enrichment of immunogenic epitopes, and subsequent identification of anti-microbial antibody and autoantibody richness and abundance. Key findings include that (1) the DM-specific IgOme was characterised by high microbial diversity, (2) antibodies against viruses were over-represented and species of the Poxviridae family were significantly enriched, (3) DM-specific accumulated autoantibodies target a significant portion of the human proteome, including proteins from specific biological processes and interferon regulated proteins, (4) autoantibodies against TRIM33 and eleven further TRIM proteins were identified, and (5) specific TRIM proteins share epitope homology with viral species identified from the plasma of DM patients.

The identified microbial epitopes in both DM and healthy controls (HC) mapped to bacteria, viruses, fungi and archaea demonstrating the human capacity for building protection against a diverse group of species. The presence of a stably-enriched microbial component in DM was identified, characterised by a higher number of epitopes per species and against a wider microbial repertoire. The effect of sample size in our measurements suggests increased microbial exposure and interpersonal variability in the DM compared to the healthy group, leading to expansion of the identified DM-specific microbial signature upon screening additional samples. Even though our observations provide a static snapshot of the IgOme, the immune system has a memory and the accumulation of antibodies takes place from birth over the entire lifetime of the individual. Thus, our data suggest that DM as a clinical entity is characterised by diverse microbial exposure.

We observed that viruses are over-represented amongst the microbial species with the highest measured abundance in DM and HC. Due to the competitive design of the epitope enrichment process this observation suggests that viral exposure has a significant role in DM. We observed enrichment of the Coronaviridae, Herpesviridae, Orthomyxoviridae, Geminiviridae and Poxviridae families in dermatomyositis. Coronaviridae was the most enriched consisting of primarily SARS coronaviruses, Orthomyxoviridae contained primarily Influenza A viruses, Herpesviridae included A and B groups, various Geminiviridae plant viruses probably due to immune exposure via the gastrointestinal tract, and finally the Poxviridae Smallpox (Variola) and vaccinia viruses. Notably, epitopes against the Poxviridae species had the highest abundance. The smallpox virus Variola was eradicated by introduction of the vaccinia virus (VACV)-based vaccine (Sanchez-Sampedro et al., 2015). The last world-wide vaccination program was in 1967(Sanchez-Sampedro et al., 2015), and smallpox immunity persists for up to 88 years (Taub et al., 2008), therefore given the age of our donors, it is very likely that the DM associated antibodies to Poxviridae were raised through the vaccination process. This is supported by the fact that VACV antibodies were observed in the healthy sample without being enriched. Many other viral species capable of directly infecting muscle tissue were enriched in DM including Human immunodeficiency virus 1 (HIV), multiple Human papillomavirus virus (HPV) strains, Hepatitis C and B (HBV), Enterovirus A71 and Foot-and-mouth disease virus, Human parvovirus B19, and Human Adenovirus and Rotaviruses which have been identified in sporadic cases of infectious myositis(Crum-Cianflone, 2008, 2010; Xiu et al., 2013). DM plasma also was enriched in viruses reported to affect the musculature indirectly via immune mechanisms, such as Influenza, Enteroviruses, HIV, SARS-Coronavirus, Herpes viruses and Parvovirus B19, and which have been implicated in the pathogenesis of polymyositis and DM (Gan and Miller, 2011; Marie et al., 2005). Interestingly, there are case studies which report development of DM after vaccination against some of these viruses, notably HBV, HPV, influenza and VACV(Ignasi Rodriguez-Pinto, 2015). Overall, the DM-specific viral signature includes viruses which can either directly infect muscle tissue and/or indirectly sabotage immune homeostasis. The observation that the respiratory and gastrointestinal systems are the primary physiological targets of these viruses agrees with previous observations regarding the high frequency of this type of infection in patients with juvenile- and adult-onset DM (Pachman et al., 2005; Svensson et al., 2017). Moreover, the age-range of juvenile and adult onset DM coincides with the epidemiological peaks of respiratory infections during life(Kuan et al., 2019).

In DM, we identified autoantibodies against a very large number of human proteins (>2500), representing about 14% of the current human proteome (UniProtKB/Swiss-Prot database, last modified 29 July 2019). This was double the proportion compared to the HC-enriched proteome, with only 13% of autoantibodies being found in both DM and healthy controls, suggesting accumulation of autoantibodies in DM against a wider group of proteins. Interestingly three times the number of autoantibody targeted proteins were regulated by type I and II interferons in DM than in HC. This underscores the relevance of interferons in this disease(Neely, 2019; Somani et al., 2008), and indicates the potential of interferon blockade as a therapy. The identification with high confidence of specific biological processes enriched in both DM and HC, suggests an organisational structure within the targets of these accumulated autoantibodies. Although deconvolution to individual samples is not possible with our data, this is consistent with an infection (exposure)-driven pathological autoantibody-producing mechanism which progressively expands its repertoire (antigens) under the chronic burden of accumulated stimuli(Chan and Gack, 2016; Cusick et al., 2012; Munz et al., 2009; Panoutsakopoulou et al., 2001). Biological processes enriched in DM are orientated around cytoskeletal organisation, microtubule movement, actin filaments and cell junctions. Eight of the identified protein targets participated in most of these processes indicating a key role in the regulation of multiple signalling pathways including the formation and disassembly of focal adhesions. The enrichment of antibodies against proteins with a role in the regulation of focal cell-cell adhesion in DM is particularly interesting, since disruption of these structures in epithelial cells is known to facilitate the spread of viruses, by allowing their release from the basal to the apical surface or the external environment (Spear, 2002; Torres-Flores and Arias, 2015). Interestingly some of the DM associated viruses described here are known to manipulate junctional proteins namely Adenoviridae, Flaviridae, Herpesviridae, Retroviridae, Paramyxoviridae, and Picornaviridae which provides a functional link between DM and increased viral exposure(Mateo et al., 2015; Mothes et al., 2010; Zhong et al., 2013).

We identified autoantibodies against TRIM33 in DM plasma, confirming the anti-TIF1 positive profile of our patients. Furthermore, we identified autoantibodies against 11 other TRIM proteins in DM, of which only TRIM21 was identified in healthy controls but with much lower abundance. TRIM21 (Ro52) is a known autoantigen in IIM and related autoimmune disorders, and acts as the highest-affinity Fc receptor in humans. TRIM21 binds cytosolic antibodies bound to non-enveloped viral pathogens, to trigger antibody-dependent intracellular neutralization and proteasomal degradation of the immune complex (Lee, 2017; Rajsbaum et al., 2014). The autoantibodies we identified against TRIM proteins, other than TRIM33 and TRIM21, have not been observed previously in IIM. TRIM69 inhibits vesicular stomatitis virus transcription and interacts with dengue virus non-structural protein 3 to mediate its poly-ubiquitination and degradation (Kueck et al., 2019; Wang et al., 2018). TRIMs also regulate antiviral pathways indirectly by mediating innate immunity, influencing the transcription of TI-IFNs, pro-inflammatory cytokines and interferon-stimulated genes (van Tol et al., 2017). Notably, the TRIM protein family expanded very rapidly in evolution, coinciding with development of the adaptive immune system, suggesting that TRIMs may have evolved to fine-tune interactions between the increasingly complex innate and adaptive immune systems (Rajsbaum et al., 2014; Versteeg et al., 2014). Deregulated activity of approximately one third of the TRIM proteins has been associated with the development of various type of human cancer (Vunjak and Versteeg, 2019). For example, TRIM47 and TRIM27 have an oncogenic role in colorectal, prostate, oesophageal, ovarian and non-small cell lung cancer, promoting proliferation and metastasis(Han et al., 2017; Liang et al., 2019; Zhang et al., 2018). Conversely TRIM3 has tumour suppressor activity, inhibiting the growth of liver, colorectal and gastric cancer(Boulay et al., 2009). The presence of autoantibodies against cancer-associated TRIMs accords with the strong temporal association between myositis and the development of malignancies in adult-onset anti-TIF1γ positive DM(Oldroyd et al., 2019), consistent with the high proportion of cancer-associated myositis cases in the current study (45%). Overall, the expanded targeting of TRIM proteins we observed in DM plasma supports the role of the TRIM RING-type E3 ubiquitin ligase subfamily as powerful regulators of the immune system through post-translational modification (Hage and Rajsbaum, 2019; van Tol et al., 2017).

Since Poxviridae epitopes were enriched in DM, we investigated a potential link between pox viruses and TRIM proteins. We identified epitope sequences with high similarity between TRIM3 and Variola virus in addition to other members of the Poxviridae family. Moreover, TRIM47 had epitopes in common with HIV1 and Synechococcus phage. These high similarity epitopes were identified in the C-terminus of the TRIM proteins; between the filamin and NHL 1 domains for TRIM3, and proximal to the B30.2 domain in TRIM47. The C-terminus of TRIM proteins is under evolutionary positive selection and determines ligand binding specificity, function and subcellular localization; notably the B30.2 domain is associated with ability to restrict viral infection, particularly retroviruses (van Tol et al., 2017). This finding of high similarity epitope sequences suggests that molecular mimicry, at least amongst specific viral and TRIM epitopes, is a potential mechanism of pathogenesis in DM.

The finding of antibodies against a human protein does not mean that this was the antigen against which the antibody was raised, as pathogen-derived proteins can mimic endogenous epitopes(Panoutsakopoulou et al., 2001; Walker and Jeffrey, 1986). This might explain inconsistencies between the universal cellular distribution of specific antigens and the specific skeletal muscle target of the disease(Walker and Jeffrey, 1986). In support of this, we observed that certain DM-specific TRIM epitopes are shared amongst different viruses. TRIM epitopes did not form a separate phylogenetic cluster but were dispersed throughout the viral epitope phylogenetic tree, suggesting that accumulated antibodies against TRIM proteins might be the product of a primal molecular mimicry event and subsequent epitope spreading rather than increased TRIM protein production. This is supported by the observation that the gene expression levels of autoantigens in myositis muscle biopsies do not correlate with the levels of circulating autoantibodies recognizing each cognate endogenous autoantigen; instead they are directly associated with the expression of muscle regeneration markers(Pinal-Fernandez et al., 2019).

We propose a model whereby pathogenesis of DM is dependent on dynamic and personalised viral exposure patterns, which start very early in life. The main two gateways of physiological transmission of viruses to humans are the respiratory and gastrointestinal systems, while non-physiological presentation of viruses is through immunisation. The link between DM and viruses is complex but strikingly present in our data since there is systematic enrichment for human proteome modules that are directly exploited by, or respond to, most of the viruses that we identified. We propose that an initial viral infection or virus delivery system presents the main antigen that induces antigen-specific antibody production in a predisposed immune environment. Since TRIM proteins share epitope homology with multiple viral species, these antibodies can also bind to TRIMs and potentially other homologous proteins. The outcome of this process is influenced by factors including the antibody populations at the site of infection (local antibodies), the status of the innate immune system, the virus-specific antibody concentration in serum (which usually peaks several weeks post infection), and the history of previous exposure primarily to the same or related viral strains (and epitopes)(Rojas et al., 2018). Since HLA Class II molecules are major regulators of the adaptive immune response to antigenic challenge through T cell repertoire selection, the level of antibody production to different viral antigens also may be mediated by HLA type. We previously identified that adult-onset anti-TIF1 positive DM is associated with HLA-DQB1*02:02(Rothwell et al., 2019). For the samples with available data included in this study, we observed that 6/15 (40%) DM and 3/17 (18%) HC are heterozygous for *HLA-DQB1*02:02*. The finding that in myositis, autoantigen expression correlates with expression of muscle regeneration markers(Pinal-Fernandez et al., 2019) suggests that after initial viral presentation, the increased concentrations of muscle proteins may then be sufficient even in the absence of viral proteins to invoke periodic rises of autoantibodies. Moreover, the continuous activation of innate immunity observed in DM(Neely, 2019; Pinal-Fernandez et al., 2019; Walsh et al., 2007; Wong et al., 2012) can also potentiate autoimmunity through chronic immune-mediated tissue damage resulting in autoantigen release without the need for specific activation of auto-reactive T cells by a microbial mimic(Panoutsakopoulou et al., 2001). Therefore, following the initial viral exposure there is a high chance that this effect will spread amongst related proteins within specific signalling pathways (since protein homology is related to function) of the initial hit. In this model, in specific autoantibody-positive myositis subgroups, accumulated autoantibodies against proteins that participate in specific biological processes and signalling pathways would be observed. The finding that in healthy controls the protein coverage of the GO processes significantly decreases suggests more random targeting of human proteins by autoantibodies, opposite to that observed in DM and in support of our hypothesis.

The use of pooled plasma is a limitation in our study since we cannot deconvolute the data into individual patients. However, the enrichment process against healthy adults provides a unique opportunity to study DM as a disease system capturing the breadth of microbial antibody and autoantibody accumulation against a very large number of linear epitopes. Apart from antibodies against TIF1 which is a shared feature of the patients in this study, we do not anticipate that our findings will be equally distributed among patients. Indeed, if the shared feature in DM is a molecular mimicry event combined with epitope spreading, then the stochastic nature of microbial exposure, genetic predisposition and the dynamic nature of immunity would not support a single pathotype. Thus, our study provides the first DM-specific “map” of potential routes to disease. For the future, we recommend: clinical records are kept of short and long-term microbial exposure, infections history including type of pathogen, severity scores, number of hospitalisations and vaccination history; which were not available in this study. Records could be expanded in cases of juvenile DM, to include maternal infection and vaccination history to account for acquired transplacental immunity.

Lifelong exposure to viruses and probably to other microbes contributes not only to accumulation of virus-specific antibodies and protection, but generation of autoantibodies against TRIMs and other cellular proteins, which at some point may reach a critical mass and induce disease most likely as a result of seemingly mild or non-symptomatic infection. Molecular mimicry and epitope spreading may play a significant role, instantly raising questions not only concerning the extent but also the sequence of events. This also means that autoantibodies identified in DM to date might only be the tip of the iceberg.

## Acknowledgements

This study was supported by a research grant from The Myositis Association. TDJW was supported by a research grant from Children with Cancer and The Caring Cancer Trust. XH was supported by grants from The Humane Research Trust and work in the Viral Oncology Labs was supported by grants from the Cancer Prevention Research Trust. HC and JL were supported by the Medical Research Council (MR/N003322/1). HC was supported by the NIHR Biomedical Research Centre Funding Scheme. The views expressed in this publication are those of the authors and not necessarily those of the NHS, the National Institute for Health Research or the Department of Health.

## Author contributions

Conceptualization, I.H., L.H., T.D.J.W., X.H., S.M.,W.E.R.O., and J.A.L.; Methodology, I.H., L.H., T.D.J.W., X.H., S.M.; Software, T.D.J.W., X.H.; Investigation, S.M., T.D.J.W., X.H., and J. O’S.; Resources, I.H., L.H., N.P., and A.P.; Writing – Original Draft, S.M., T.D.J.W., and J.A.L.; Writing – Review & Editing; S.M., T.D.J.W., J.A.L., I.H., L.H., X.H., W.E.R.O., H.C., J.O’S., N.P., and A.P.; Visualization, S.M., T.D.J.W.; Supervision, I.H., L.H., and J.A.L.; Funding Acquisition, J.A.L., I.H., W.E.R.O., and H.C.

## STAR methods

### LEAD CONTACT AND MATERIALS AVAILABILITY

Further information and requests for resources and reagents should be directed to and will be fulfilled by the Lead Contact, Janine Lamb.

This study did not generate new unique reagents.

### EXPERIMENTAL MODEL AND SUBJECT DETAILS

#### Study cohort

Plasma samples were collected from anti-TIF1 positive adult-onset dermatomyositis (DM) patients through the UK Myositis Network, as described previously (Rothwell et al., 2019) (Table S1). All individuals fulfilled definite or probable Bohan and Peter classification criteria for dermatomyositis. Anti-TIF1 autoantibody positivity was identified by immunoprecipitation, and confirmed by ELISA, as described previously(Rothwell et al., 2019). Gender and age matched (at time of sample collection) healthy controls (HC) were identified through the University of Manchester Longitudinal Study of Cognition in Normal Healthy Old Age cohort (Rabbitt et al., 2004). All samples were collected with relevant research ethics committee approval (MREC 98//8/86 North West Haydock Research Ethics Committee for UKMyoNet and UREC 02225 and UREC4 2017-1256-2489 for healthy control cohort). Study participants provided written informed consent. Clinical, demographic and experimental information are presented Table S1.

### METHOD DETAILS

#### Experimental protocol

We implemented the “Serum Antibody Repertoire Analysis (SARA)” pipeline (manuscript in preparation). SARA comprises a comprehensive workflow that integrates molecular biology peptide display and epitope signature enrichment through competitive bio-panning and high-throughput DNA next generation sequencing (NGS), alongside in-house computational scripts to reverse engineer high resolution epitope signatures that reflect original *in vivo* epitopes present in patient sera. SARA was applied to predict the identity and abundance of antibody epitope repertoires enriched in plasma from anti-TIF1 autoantibody-positive DM patients versus plasma from matched HC (see Supplementary Methods). This pipeline provides a digital triage of infectious organism epitopes and autoantibodies predicted to be uniquely present or highly enriched in each sera pool. All informatics analysis was carried out using R(R Development Core Team, 2018) 3.2-3.6 and Python(Sanner, 1999) 3.5. We briefly summarise the seven SARA pipeline modules (M1-M7) below in turn.

#### Serum Antibody Repertoire Analysis pipeline

##### Plasma total immunoglobulin purification: M1

Total immunoglobulins IgA, IgG, IgM were purified from twenty anti-TIF1 positive adult-onset DM and twenty HC plasma by AdamTech Total Ig Extraction kits (Pesac, FR). For the sample-pools of 10 (P10), 12 µg of isolated Ig per donor were used for a total of 120 µg. For the sample-pools of 20 (P20), 6 µg of isolated Ig per donor were used for a total of 120 µg. Ig yields and purity were assessed by NanoDrop spectrophotometry and polyacrylamide gel electrophoresis.

##### Competitive biopanning: M2

Separate purified Ig pools from DM and HC were used for competitive bio-panning with the FliTrx^TM^ random 12 amino acid (AA) peptide surface display system(Lu et al., 2003). Briefly, we first conducted a pre-panning stage to incubate separate DM and HC immobilised Ig pools with induced FliTrx^TM^ *E. Coli* cells(Lu et al., 2003) in order to sequester expressed epitopes relevant to the specific plasma cohort. Unbound bacteria per cohort were further incubated with the alternative immobilised Ig pools for the main panning stages (cross-panning). Tethered bacteria were eluted and expanded to repeat the biopanning process 5 times. OD600 was measured after each round of biopanning to ensure comparable efficiency between DM and HC (Figure S3E). After competitive biopanning immobilised bacteria were expanded and polyvalent plasmid cohorts were purified (Maxiprep, Qiagen, UK).

##### Epitope NGS: M3, NGS data processing: M4

Polyvalent plasmid variance regions were PCR amplified and gel-purified. PCR products were validated by Sanger sequencing (Figure S3F), and NanoDrop. The DNA primer sequences are: FliTrx^TM^ Forward: 5’-ATTCACCTGACTGACGAC-3’, FliTrx^TM^ Reverse: 5’-CCCTGATATTCGTCAGCG-3’. Multiplexed DNA fragments were sequenced on the NextSeq 500 platform (Illumina, UK). We retrieved ≈24 million and ≈36 million paired end FASTQ reads for HC and DM respectively (Figure 1). The 36-bp variance region sequences were translated with respect to reading frame. < 14% of DM and <13% of HC comprised premature stop codons, indicative of immunologically-relevant bio-panning enrichment (Figure S1A-S1D). Non-specific HTS sequence noise from residual bio-panning solution was controlled for by our 10-σ-99 noise floor. This retains all expressed epitopes within each respective HC or DM pool that surpass a threshold of 10 standard deviations above the lowest 99% distinct peptide sequence enrichment when ranked by read count (further described in Supplementary figure S1A-S1D).

##### Epitope signature set analysis: M5

Distinct epitope signature set analysis presented unique and common AA epitope sequence sets with associated NGS enrichment scores. Minimum thresholds of 10AA sequence length and the 10-σ-99 NGS noise filter provided sensitive annotation power while minimising annotation noise (Figure S1D-S1F). Cleaned distinct epitope sequences were collated. Unique epitope pools were determined by symmetric difference of the HC and DM pools (i.e. specific epitope sequences uniquely found in HC or in DM patients). The intersect between HC and DM pools was probed and a minimum five-fold change threshold applied to partition relevant epitope collections to supplement the HC or DM pools. These epitope collections were utilised in downstream microbiome and autoantibody annotations. >8.6×10^6^, and > 4.7×10^6^ distinct epitopes with a minimum length of 6AA length were retrieved for DM and HC pools respectively. The highest NGS read counts achieved were ≈111,000 (DM) and ≈88,000 (HC) (Figure 1).

##### Epitope annotation and rank scoring: M6

15,522 DM-associated and 4,817 HC-associated unique epitopes were retrieved for DM patients. Only 0.65% of the DM pool comprised peptide sequences common to HC. Of these epitopes common by sequence, the DM enrichment score ranged between 5-fold and 546-fold the healthy control score (Figure 1) (Figure S1A-S1F). Epitopes were annotated by our modified BLASTp(Henikoff and Henikoff, 1992) approach at 100% minimum identity (Figure S1G-S1I). This facilitates short AA-sequence inputs and resolves protein annotation confidence scores per epitope with statistical control of epitope-protein annotation confidence, intra-protein dis-contiguous sequence matches, redundancy of protein isoforms and common sequence matches, single-epitope multi-protein parsing, and set analysis of resultant patient cohort organisms lists. These were tuned per each epitope’s NGS read count. 6.745×10^6^ (DM patients) and 2.250×10^6^ (HC) total protein annotations were retrieved (Figure 1).

##### Phylogenetic and taxonomic analysis: M7

Our scoring system generates aggregate NGS read count enrichment and annotation stringency scores. We retrieved every known upstream taxonomic ranking level for all microbe IDs and anchored these via NCBI REST and the Taxonomy repository (Benson et al., 2009; E., 2008; Federhen, 2012). Phylogenetic trees were generated by PhyloT biobtye(Letunic, 2018) and microbial annotations were assessed by NGS enrichment, epitope mapping confidence, annotation scores, taxonomic ranks, and phylogenetic clustering (Figure S1G-S1I). We utilised the interactive Tree Of Life iTOL(Letunic and Bork, 2007; Letunic and Bork, 2016) to taxonomically cluster microbial agents for the DM or HC pools with incorporated annotation layers representing the NGS enrichment and annotation scores (Figure S1I). 9,111 distinct and 6,202 unique species were annotated for DM patients (4,994 distinct and 2,085 species were annotated for HC) (Figure 1).

#### Human protein autoantibody digital triage

We identified any autoantibodies against human proteins that may be present. Annotation scores occupy unit interval space [0,1] as surrogate significance values. Protein cohorts were inspected by Metscape(Tripathi et al., 2015), KEGG(Kanehisa and Goto, 2000), UniProt(Apweiler et al., 2013), and the Human Protein Atlas(Uhlen et al., 2010), to explore functional predictions based on aberrant autoimmune targeting. 435 (DM) and 407 (HC) enriched human biomarkers from autoantibodies were retrieved (Figure 1).

#### Phylogenetic analysis of epitope sequences

Alignments of human and viral epitope sequences were performed with a gap open cost of 10.0 and a gap extension cost of 1.0. Tree (circular and radial) construction was performed using Neighbour joining and Kimura protein distance measure with bootstrap analysis (100 replicates). Analysis was performed in CLC Genomics workbench 12.

### QUANTIFICATION AND STATISTICAL ANALYSIS

Continuous variables were log10 transformed and tested for normal distribution using the Shapiro-Wilk test. Pairwise quantitative differences between two groups were tested using two-tailed Mann Whitney (Wilcoxon rank sum test) for variables not following a normal distribution and unpaired t test for variables with a normal distribution. F test was used to test for equal variance. The Kruskal-Wallis nonparametric test was used to compare three or more groups; Dunn’s multiple comparisons test was used to compare pairwise differences in the sum of ranks with the expected average difference (based on the number of groups and sample size per group). Z score transformation was used to specify the location of each observation within a distribution. Statistical analyses were performed with GraphPad Prism 8.0 (GraphPad Software, Inc., San Diego, CA)(GraphPad Software).

### DATA AND CODE AVAILABILITY

The code supporting the current study have not been deposited in a public repository but will be deposited following submission of our methods manuscript (in preparation). In the intervening period these will be available from the corresponding author on request.

#### Supplemental information

Supplemental information contains Supplementary methods, Supplementary figures 1-7, and Supplementary tables 1-3.

#### Supplementary Methods

##### Serum Epitope Repertoire Analysis (SARA) of DM patients

DM total Ig purification (M1); Competitive biopanning (M2); NGS epitope sequencing (M3). We retrieved ≈24 million and ≈36 million paired end FASTQ reads for HC individuals and DM patients respectively (Fig. S1A). These were highly enriched by associated NGS read counts and represent polyvalent FliTrxTM peptide 12 AA coding sequences from DM and HC biopanned total Ig pools.

##### DM NGS data processing (M4); DM epitope signature set analysis (M5)

We retrieved 13,385,371 distinct expressed AA epitopes that represent the epitope signatures present within the DM and HC Ig pools (Fig. S1A). Associated NGS read frequencies reflected the epitope enrichment process achieved during biopanning, and Fig. S1B & C confirms that the 10-σ-99 NGS filter optimally controlled biopanning-induced sequencing noise to successfully provide 15,522 DM-associated and 4,817 HC-associated unique epitope sequences. Preferential enrichment for epitopes of length 11 and 12 AAs (Fig. S1D) confirmed genuine biopanning capture and is concordant with published peptide epitope ranges of 4 to 12 AAs (Buus et al., 2012; Hopp and Woods, 1981). Minimum epitope lengths of 9AA applied with the 10-σ-99 NGS filter (above) proved optimal for annotation (Fig. S1E) using the unique sequence epitope sets of 15,522 for DM and 4,817 for HC individuals (Fig. S1F). Ig repertoire sequence overlap determined that only 0.65% of DM epitope set were present in healthy controls (fold-change (FC) values of >=5x to 546x abundance vs HC individuals) while 6.44% of healthy epitope set was retained in the DM group (FC values of >=5.3x to 1240x vs DM) (Fig. S1F).

##### DM epitope annotation (M6) with phylogenetic and taxonomic interface (M7)

Our modified BLASTp approach retrieved 6.75 million (DM) and 2.25 million (HC) microbial or human protein annotations respectively which mapped to ≈14,000 microbial organisms. Enrichment plots revealed 4,994 Distinct infectious agents in HC individuals and 9,111 infectious agents in DM patients (Fig. S1G) of which 2,085 and 6,202 Unique infectious agents were identified in healthy individual and DM patients respectively (Fig. S1H). These signatures presented 377 highly enriched infectious agents in DM and 270 highly enriched agents in healthy individuals ranked by phylogeny (Fig. S1I).

## Supplementary Figures

**Supplementary figure 1:**
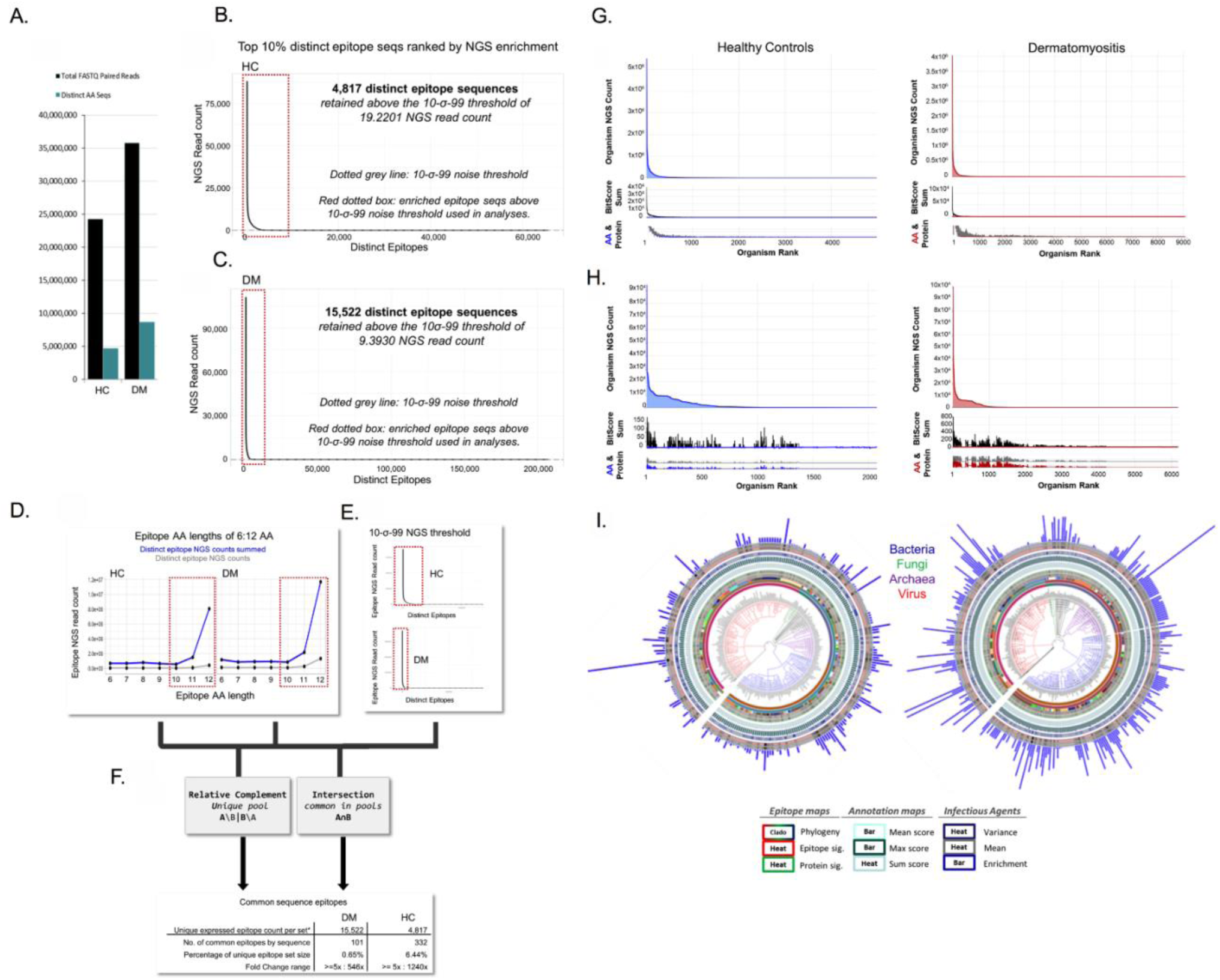
Immunoglobulin epitope enrichment in DM and HC. Next generation sequencing run data processing: (A) Total NGS reads and expressed amino acid (AA) sequences per healthy control (HC) and dermatomyositis (DM) samples. Distinct epitope sequences are ranked by their NGS read count in (B) HC, and (C) DM patients; Graphs depict the top 10% ranked epitopes per pools. The 10-σ-99 noise-floor threshold is derived from 10 standard deviations of the lowest 99% ranked epitope read counts within each pool. This was used to prevent non-specific residual biopanning peptides sequences from entering the SARA annotation module M6 (M6: Epitope annotation) (Supplementary Methods). Red dotted boxes thus represent the proportion of clean epitope AA sequence data carried forward to annotation. DM patients produced greater than three-fold enriched distinct epitope sequences vs HC (15,522 epitopes in DM vs 4,817 in HC). Epitope signature set analysis: (D) Epitope enrichment in HC and DM patients stratified by AA Seq length. Blue: Summed NGS read counts per distinct AA epitope sequence. Grey: Number of distinct AA epitope sequences. Preferential enrichment for epitopes between 10 to 12 AA is clearly visible indicative of genuine antibody-epitope binding during the biopanning process. Furthermore, the DM pool preferentially enriched a larger cohort of disease-related epitope sequences compared to healthy controls. (E) NGS noise floor control (10 SD above the lowest 99% amino acid NGS read count values. Red boxes are parameters used downstream. (F) Relative complement and intersection sets. Only 101 (DM) and 332 (HC) distinct epitopes remained common to both pools after NGS fold change control. The enrichment fold-change differences of these common sequences ranged from 5x:1,240x compared with the opposite set (0.65% and 6.44% the proportion of the respective unique epitope sets). Enriched Distinct and Unique microbial agents in DM: Distinct and unique organisms at a 100% BLAST annotation threshold. Epitope sets comprised 10-12 AA sequence lengths following the 10-σ-99 NGS noise threshold. The DM and HC microbes were annotated from 6,745,487 and 2,250,772 protein matches respectively. (G) Images of interactive enrichment plots of highest ranked 4,994 distinct infectious agents (HC; left panel) and 9,111 infectious agents (DM; right panel). Top plot provides NGS enrichment score per agent; middle plot is the annotation confidence score; lower graphs are traces for number of epitopes per agent and number of proteins per agent. (H) Corresponding enrichment plots for 2,085 (HC) and 6,202 (DM) Unique infectious agents. (I) Images of interactive digital triage plots. 377 highly enriched infectious agents mapped to DM and 270 highly enriched agents mapped to HC from ≈2.25 million & ≈ 6.75 million distinct epitopes. Cladogram leaf colour keys are provided. Inner two rings: heat strips of trace data for number of epitopes per agent / number of proteins per agent. Subsequent 3 rings: annotation confidence score. Outermost three rings: NGS enrichment data per infectious agent of mean NGS counts per agent, NGS variance within each agent, and final enrichment score (outer blue bars).

**Supplementary figure 2:**
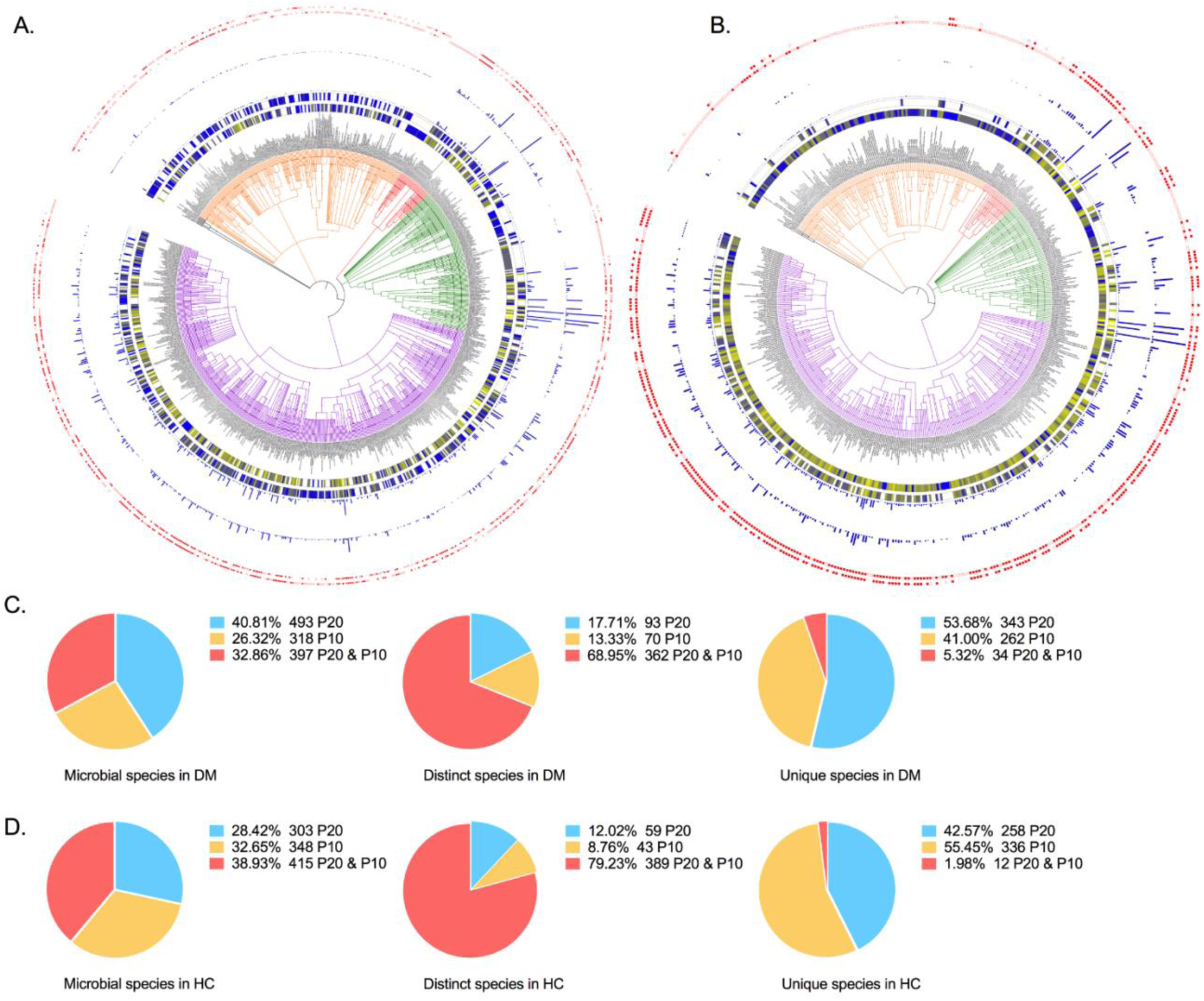
Microbial species identified in DM and HC. Taxonomic clustering of identified microbial lineages up to the species level in (A) DM and (B) HC. The microbial cladograms contain taxa ids after integration of P10 and P20 experiments. Inner ring: Microbial NGS reads per taxon identified in P20. Second ring: Microbial NGS reads per taxon identified in P10; Colour-gradient: Yellow; highest amount, Blue; lowest amount. Microbial NGS reads are log10 transformed. Bar plots represent number of different AA peptide sequences per microbial taxon. Inner ring; P20, Outer ring: P10. Squares depict whether a microbial taxon was detected in the paired DM and/or HC: No filling; unique species. Red filling; distinct species. Squares annotated in both P20 and P10 represent a stably enriched microbial component in DM or HC. Inner ring; P20, outer ring: P10. Dendogram colours: Pink: bacteria, Orange: viruses, Green: eukaryotes, Red: Archaea. Distribution of shared microbial species between the P20 and P10 pools in DM and HC: In the pie chart panel the distribution of microbial species is presented in DM (C) and HC (D) pools. In dermatomyositis 32.86% of the total identified species was shared between DM P20 and DM P10: Of the distinct species 68.95% was shared between DM P20 and DM 10; of the unique species 5.32% was shared between DM P20 and DM P10 and 53.68% was observed in only DM P20 compared to 41% observed only in DM P10. In the healthy group, 38.93% of total species was shared between HC P20 and HC P10: 79.23% of distinct species was shared between HC P20 and HC P10; 1.98% of unique species was shared between HC P20 and HC P10, and 42.57% of species was identified only in HC P20 compared to 55.45% in HC P10.

**Supplementary figure 3:**
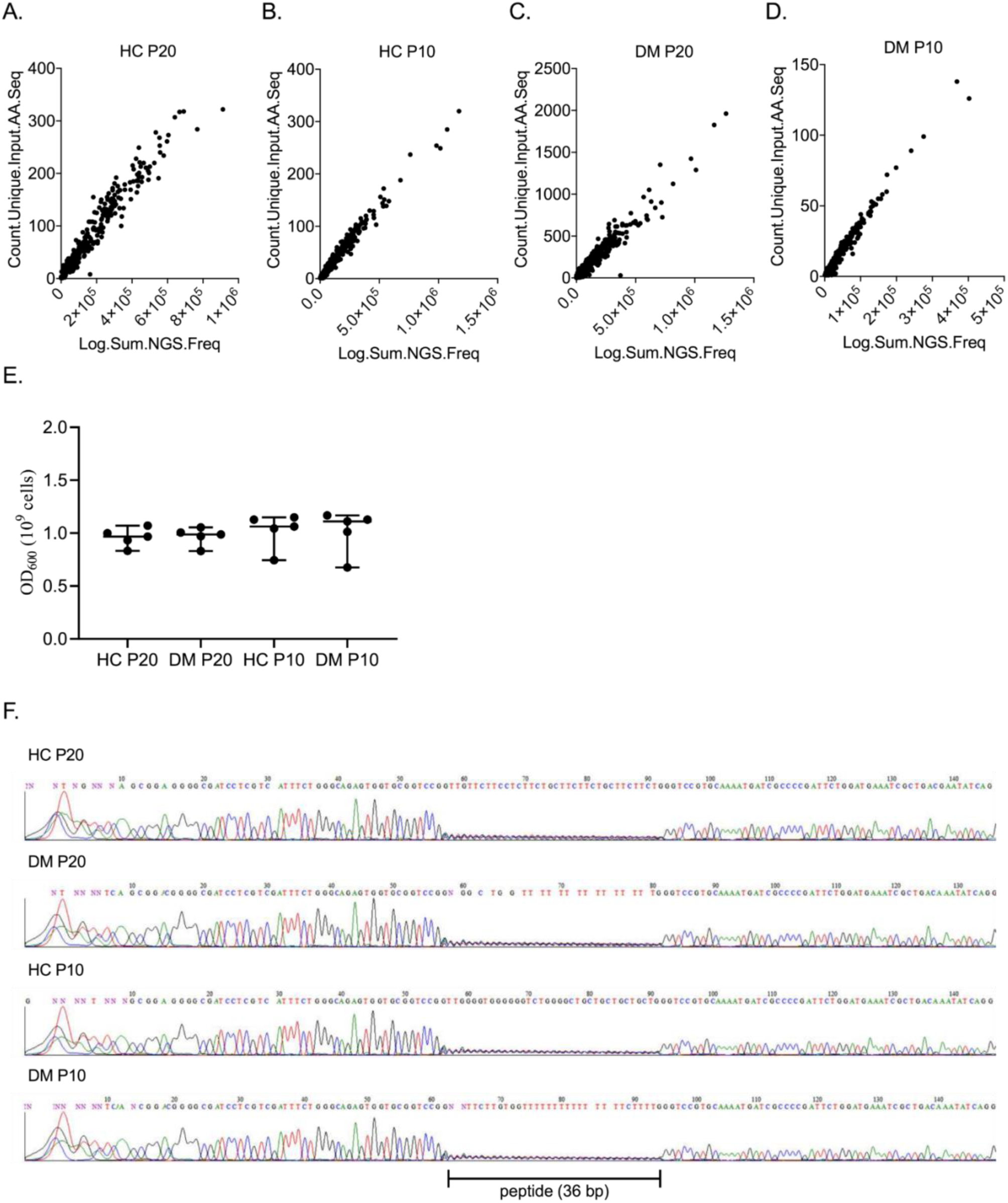
NGS data normalisation and verification of random peptide enrichment. Correlation of number of NGS reads and total number of epitopes per microbial species for (A) HC P20, (B) HC P10, (C) DM P20, and (D) DM P10.The scatter plots describe the positive and linear correlation of the sequencing depth and the number of identified microbial epitopes. To integrate both factors in the meta-analysis we defined NGSRe-norm. (E) Optical density measurements during the biopanning process. The optical density of the FliTrx E. Coli cells was measured after each round of panning. No differences were observed between the DM and HC cross-panning pairs. Welch’s t test p value 0.861 for P20 comparison and 0.954 for P10. (F) DNA chromatograph of amplified variance region. Illustrative variance region following Sanger sequencing. Portion of variance region is ¼ the intensities of flanking consensus along the 36bp variance region as a function of each measured base (“N”) comprising equal quantities of A, C, T, and G nucleotides.

**Supplementary figure 4:**
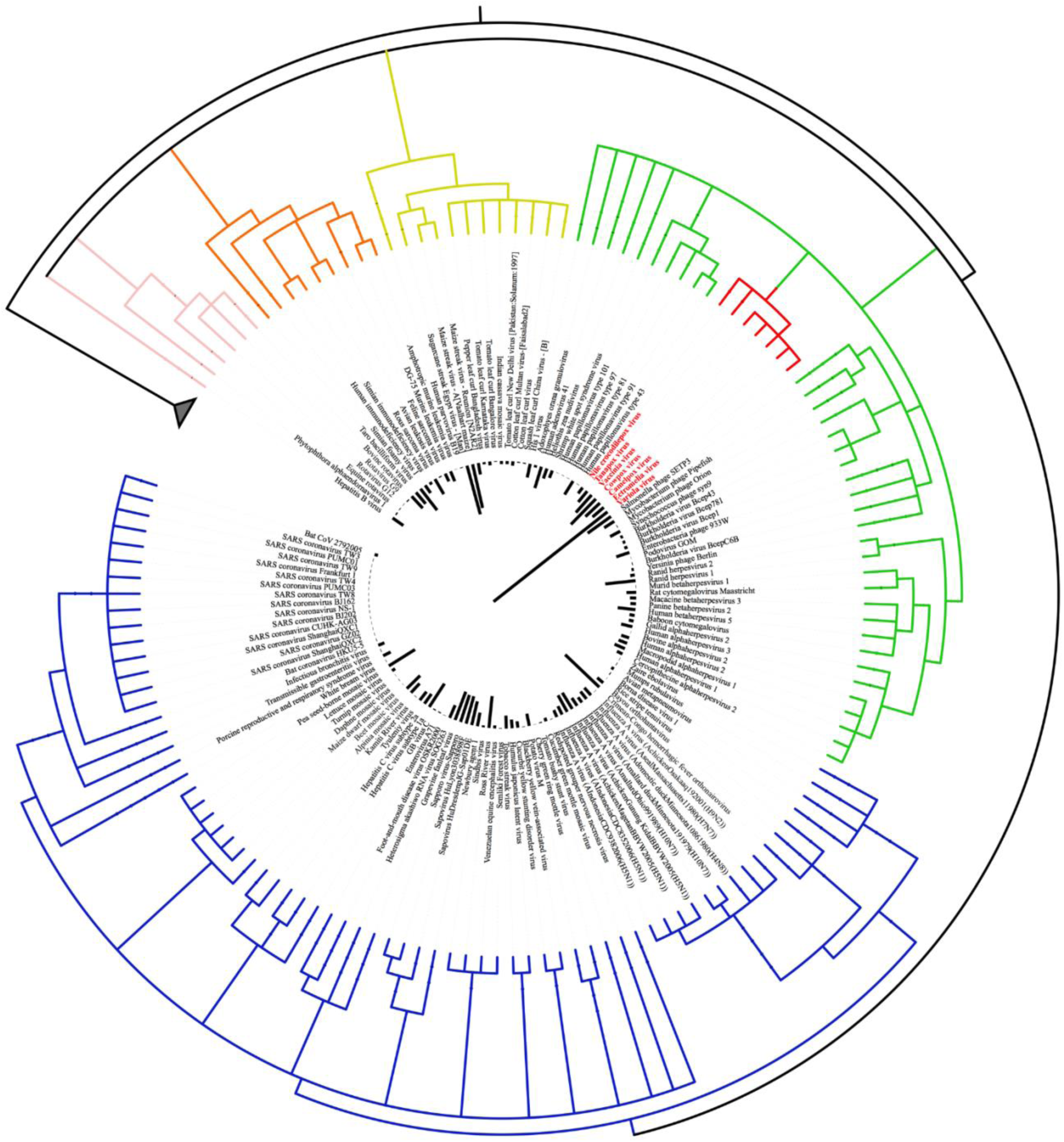
Dermatomyositis-specific viral species. Inverted circular cladogram based on the taxonomic similarity of the dermatomyositis enriched viral species in DM P20. Clade colours are representative of genome type: Blue; single stranded RNA, Green; double stranded DNA, Gold; single stranded DNA, Orange; RNA reverse transcribing, Pink; DNA reverse transcribing. Nodes with red colour depict Chordopoxvirinae (Poxviridae) species. Single bar plots (inner ring) indicate NGSRe-norm values per viral species.

**Supplementary figure 5:**
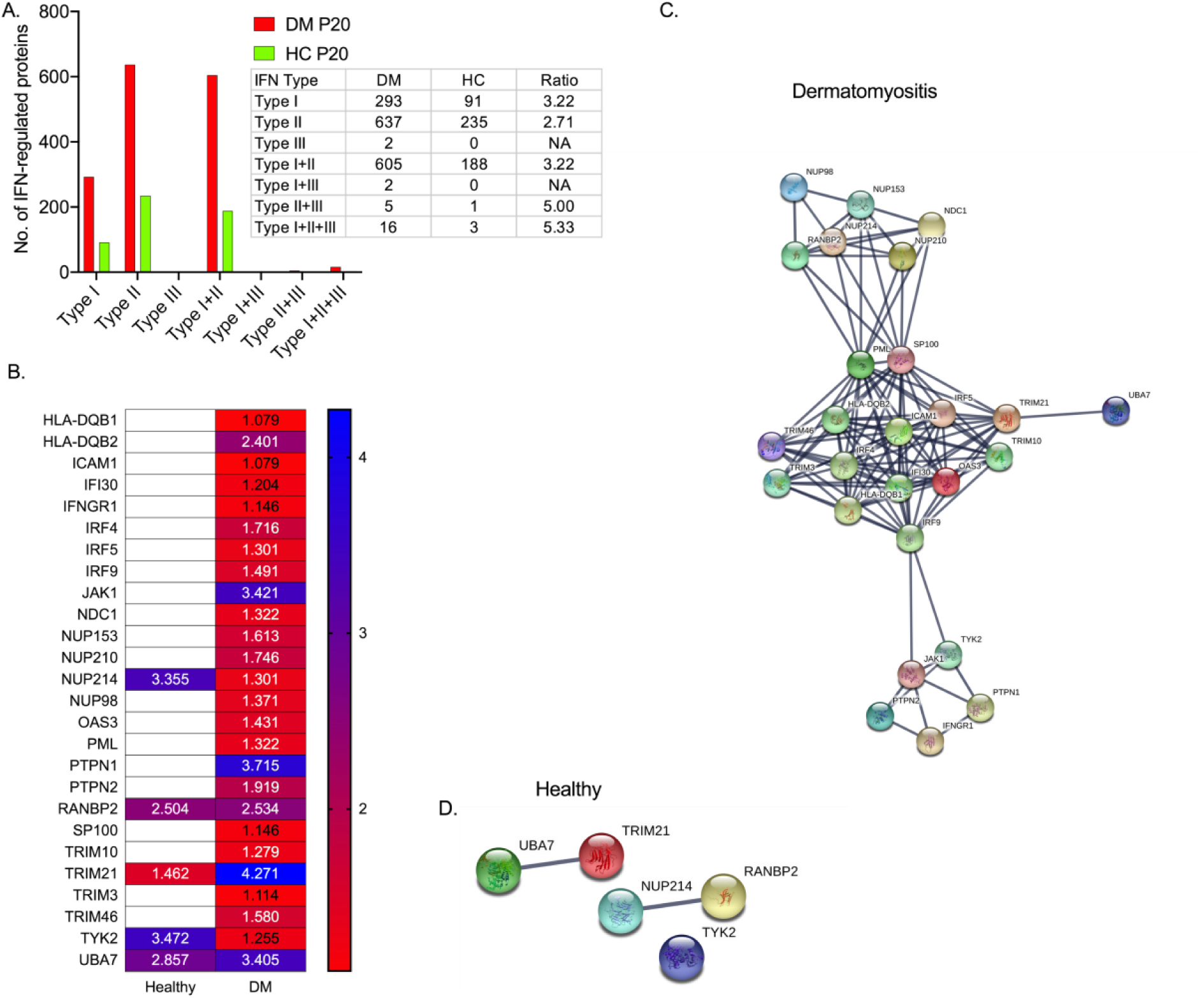
The IFN-related autoantibody proteome, focussing on IFNG. (A) Distribution of autoantibody protein-targets identified in DM and HC based on their predicted interferon-regulated profile. The majority of autoantibodies in DM target IFN-regulated proteins participating in type II alone or type I & II signalling pathways. (B) Heatmap of the autoantibody levels (mean log10 intensities) against proteins that are part of the IFNG signalling pathway in dermatomyositis and healthy controls. (C) Protein-protein interaction graph of autoantibody targets enriched in DM, and (D) enriched in the healthy sample. Each node represents an identified protein. Edges between two nodes represent confidence of interaction. Active interaction resources included experiments, gene fusion, co-occurrence, and co-expression. The minimum required score for a valid interaction was set to 0.700 (high confidence) (STRING 11.0)(Szklarczyk et al., 2019). Disconnected nodes were hidden in the networks.

**Supplementary figure 6:**
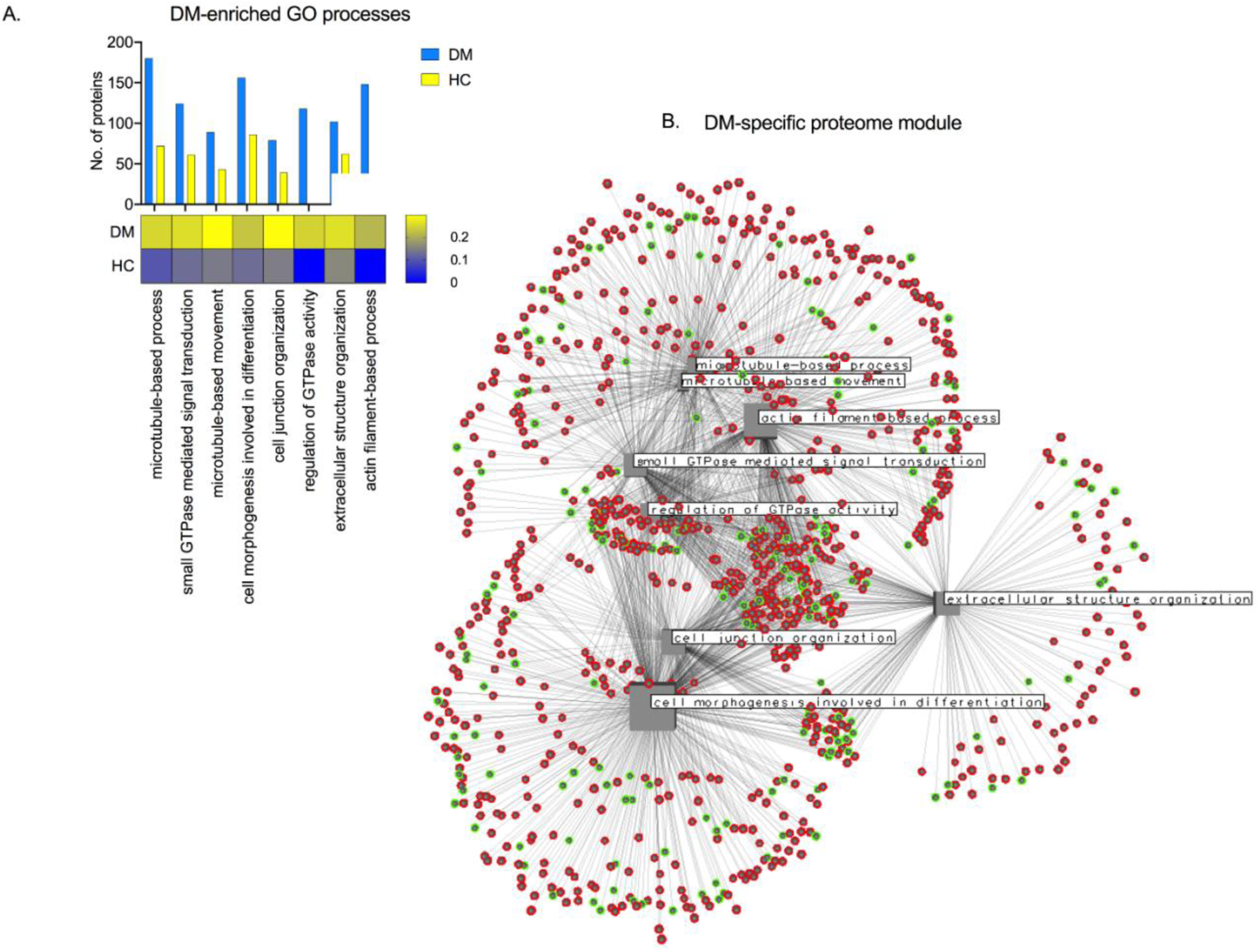
The dermatomyositis-specific autoantibody module. Human proteome autoantibody-targets that were identified in the DM P20 were used to screen for GO enrichment (biological processes). The distribution of the identified proteins in the GO processes that were successfully represented in the DM-enriched autoantibody proteome along with the GO-coverage (heatmap) in DM and HC can be seen in (A). Graph representation of the DM proteome module (B): The GO processes that were successfully represented in the DM-enriched autoantibody proteome are depicted as hubs (square nodes). Each hub is directly connected, through an edge, with the protein that is part of the specific GO process (circular nodes). Square node size is analogous to the number of interactions (edges). Nodes highlighted with red were exclusively identified in DM P20 (80% of nodes). The network contains 937 nodes (plus 8 GO hubs) with 1761 interactions. It is organised in one single component (module) with an average degree of interaction of 3.72. The eight biological functions were inter-linked because some of their protein members are shared. The network was constructed and analysed using NAViGaTOR (Network Analysis, Visualization, & Graphing TORonto)(Brown et al., 2009).

**Supplementary figure 7:**
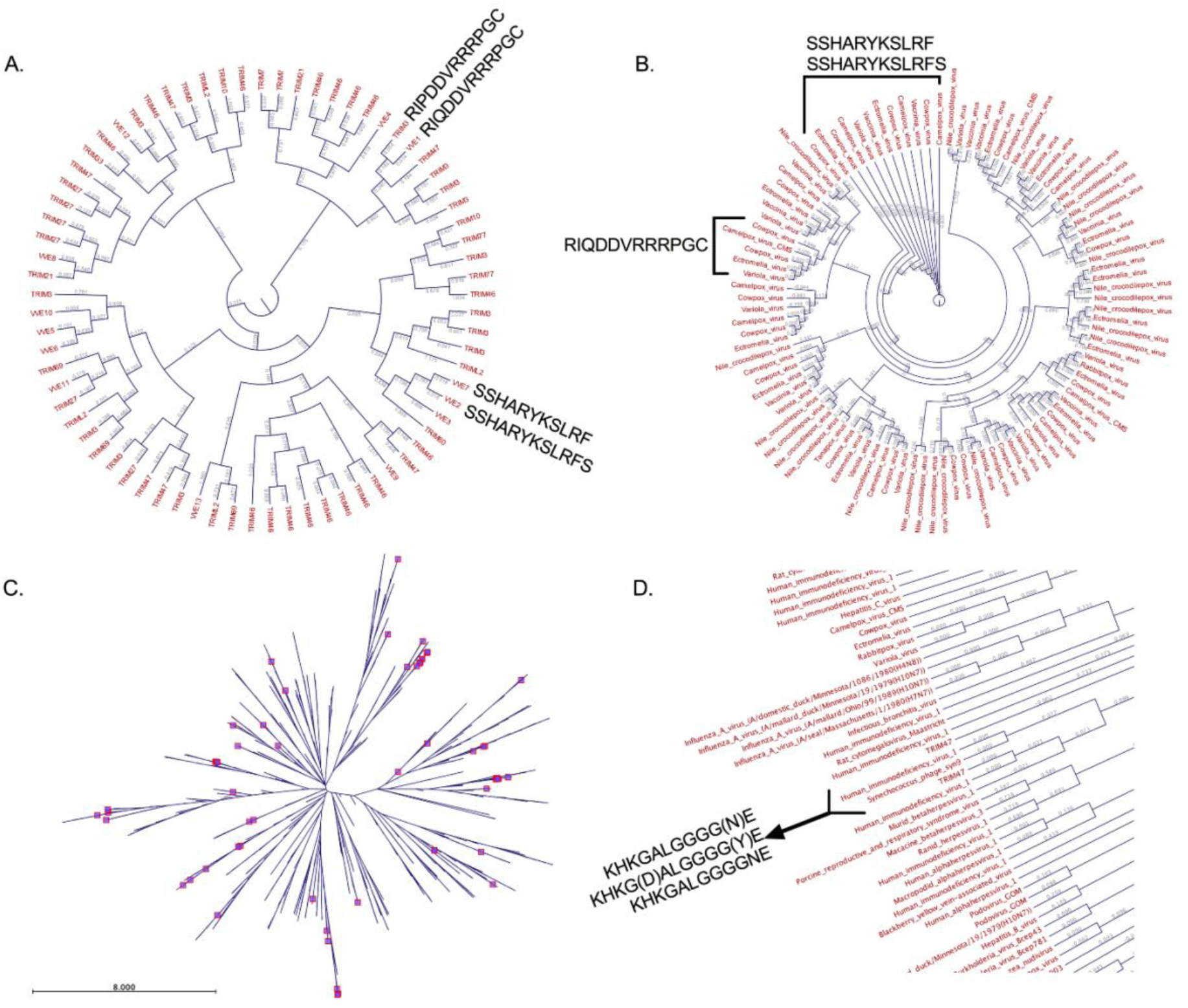
Sequence homology cladograms of epitopes identified in DM. (A) Circular cladogram of variola virus and TRIM epitopes (high specificity thresholds). VVE; variola virus epitope. (B) Circular cladogram of Poxviridae epitopes identified in DM. The three motifs which were identified to be of high similarity between variola virus and TRIM3 are also shared across the Poxviridae family. (C) Radial cladogram of aligned viral and TRIM epitopes. The TRIM epitopes are annotated with squares. Interestingly, TRIM epitopes do not form clusters and they are spread across the tree. (D) Partial circular cladogram and phylogenetic distances amongst all viral and TRIM epitopes identified in DM. The Human immunodeficiency virus 1, Synechococcus phage syn9 and TRIM47 share a common epitope (branch length=0.000). Branch lengths represent phylogenetic distance (Kimura protein distance).

## Supplementary Tables

**Supplementary Table 1:**
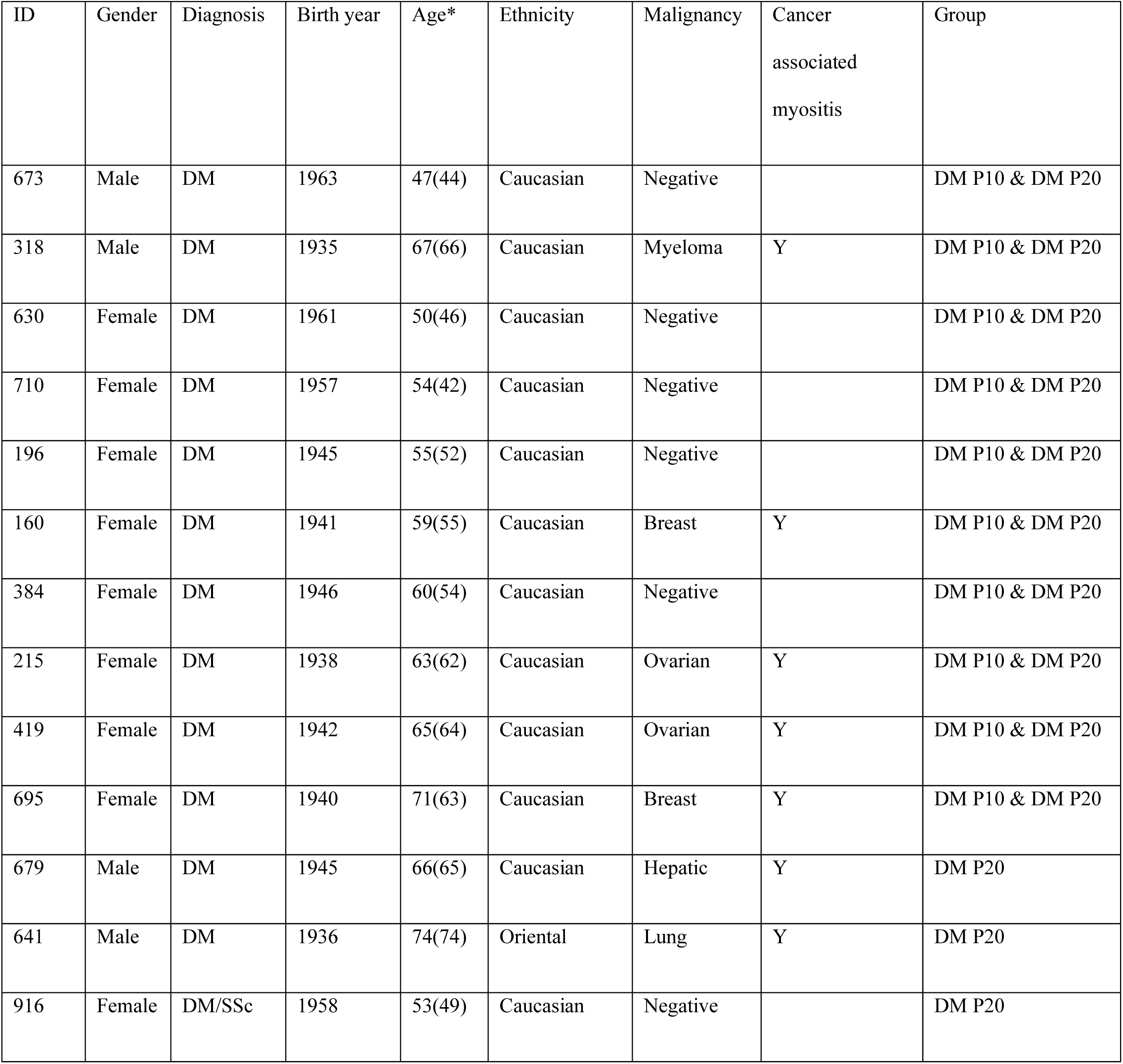
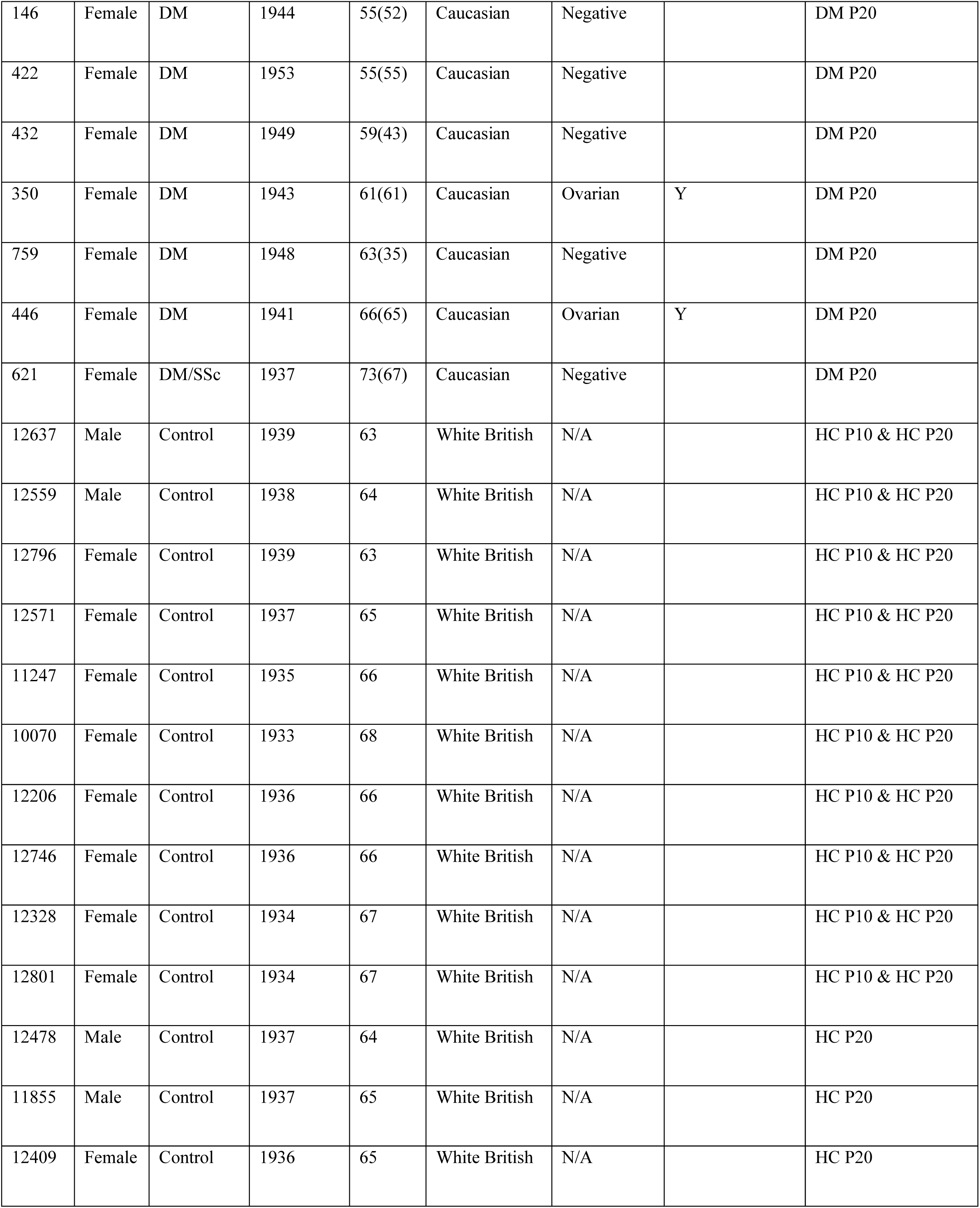
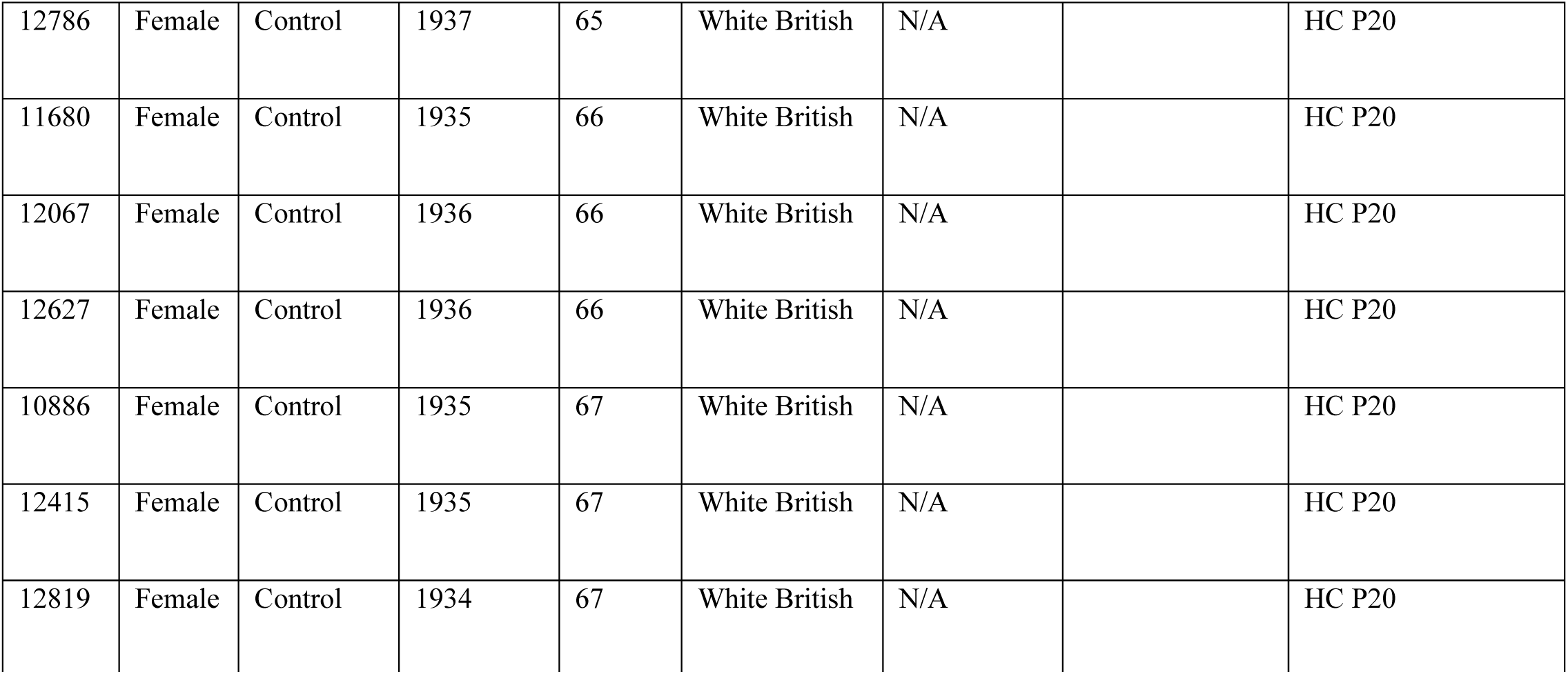
Clinical and demographic data for samples used in this study. DM; Dermatomyositis. SSc; systemic sclerosis. HC; healthy control. P10; pool of 10 samples. P20; pool of 20 samples. Cancer-associated myositis defined as cancer with three years of myositis onset. N/A; not available. *Age at time of sampling and age at DM onset.

| ID | Gender | Diagnosis | Birth year | Age* | Ethnicity | Malignancy | Cancer associated myositis | Group |
| --- | --- | --- | --- | --- | --- | --- | --- | --- |
| 673 | Male | DM | 1963 | 47(44) | Caucasian | Negative |  | DM P10 & DM P20 |
| 318 | Male | DM | 1935 | 67(66) | Caucasian | Myeloma | Y | DM P10 & DM P20 |
| 630 | Female | DM | 1961 | 50(46) | Caucasian | Negative |  | DM P10 & DM P20 |
| 710 | Female | DM | 1957 | 54(42) | Caucasian | Negative |  | DM P10 & DM P20 |
| 196 | Female | DM | 1945 | 55(52) | Caucasian | Negative |  | DM P10 & DM P20 |
| 160 | Female | DM | 1941 | 59(55) | Caucasian | Breast | Y | DM P10 & DM P20 |
| 384 | Female | DM | 1946 | 60(54) | Caucasian | Negative |  | DM P10 & DM P20 |
| 215 | Female | DM | 1938 | 63(62) | Caucasian | Ovarian | Y | DM P10 & DM P20 |
| 419 | Female | DM | 1942 | 65(64) | Caucasian | Ovarian | Y | DM P10 & DM P20 |
| 695 | Female | DM | 1940 | 71(63) | Caucasian | Breast | Y | DM P10 & DM P20 |
| 679 | Male | DM | 1945 | 66(65) | Caucasian | Hepatic | Y | DM P20 |
| 641 | Male | DM | 1936 | 74(74) | Oriental | Lung | Y | DM P20 |
| 916 | Female | DM/SSc | 1958 | 53(49) | Caucasian | Negative |  | DM P20 |
| 146 | Female | DM | 1944 | 55(52) | Caucasian | Negative |  | DM P20 |
| 422 | Female | DM | 1953 | 55(55) | Caucasian | Negative |  | DM P20 |
| 432 | Female | DM | 1949 | 59(43) | Caucasian | Negative |  | DM P20 |
| 350 | Female | DM | 1943 | 61(61) | Caucasian | Ovarian | Y | DM P20 |
| 759 | Female | DM | 1948 | 63(35) | Caucasian | Negative |  | DM P20 |
| 446 | Female | DM | 1941 | 66(65) | Caucasian | Ovarian | Y | DM P20 |
| 621 | Female | DM/SSc | 1937 | 73(67) | Caucasian | Negative |  | DM P20 |
| 12637 | Male | Control | 1939 | 63 | White British | N/A |  | HC P10 & HC P20 |
| 12559 | Male | Control | 1938 | 64 | White British | N/A |  | HC P10 & HC P20 |
| 12796 | Female | Control | 1939 | 63 | White British | N/A |  | HC P10 & HC P20 |
| 12571 | Female | Control | 1937 | 65 | White British | N/A |  | HC P10 & HC P20 |
| 11247 | Female | Control | 1935 | 66 | White British | N/A |  | HC P10 & HC P20 |
| 10070 | Female | Control | 1933 | 68 | White British | N/A |  | HC P10 & HC P20 |
| 12206 | Female | Control | 1936 | 66 | White British | N/A |  | HC P10 & HC P20 |
| 12746 | Female | Control | 1936 | 66 | White British | N/A |  | HC P10 & HC P20 |
| 12328 | Female | Control | 1934 | 67 | White British | N/A |  | HC P10 & HC P20 |
| 12801 | Female | Control | 1934 | 67 | White British | N/A |  | HC P10 & HC P20 |
| 12478 | Male | Control | 1937 | 64 | White British | N/A |  | HC P20 |
| 11855 | Male | Control | 1937 | 65 | White British | N/A |  | HC P20 |
| 12409 | Female | Control | 1936 | 65 | White British | N/A |  | HC P20 |
| 12786 | Female | Control | 1937 | 65 | White British | N/A |  | HC P20 |
| 11680 | Female | Control | 1935 | 66 | White British | N/A |  | HC P20 |
| 12067 | Female | Control | 1936 | 66 | White British | N/A |  | HC P20 |
| 12627 | Female | Control | 1936 | 66 | White British | N/A |  | HC P20 |
| 10886 | Female | Control | 1935 | 67 | White British | N/A |  | HC P20 |
| 12415 | Female | Control | 1935 | 67 | White British | N/A |  | HC P20 |
| 12819 | Female | Control | 1934 | 67 | White British | N/A |  | HC P20 |

**Supplementary Table 2:** Richness of viral families identified in Dermatomyositis and healthy controls. The identified viral families are ordered based on decreasing richness (number of species per family) in DM and HC. Colour gradient is analogous to Z-transformed values of richness (Yellow: high, Blue: low). Missing values; Non-detected viral families.

| Viral families | DM | HC |
| --- | --- | --- |
| Coronaviridae | 4.123 | 1.648 |
| Geminiviridae | 2.778 | -0.474 |
| Herpesviridae | 2.105 | 0.738 |
| Orthomyxoviridae | 1.209 | 1.648 |
| Poxviridae | 0.984 | -0.171 |
| Potyviridae | 0.760 | 0.132 |
| Flaviviridae | 0.536 | 1.648 |
| Papillomaviridae | 0.312 | -0.171 |
| Reoviridae | 0.312 | 0.738 |
| Caliciviridae | 0.087 | 2.254 |
| Podoviridae | 0.087 | -0.777 |
| Prokaryotic | 0.087 | -0.474 |
| Retroviridae | 0.087 | 1.345 |
| Togaviridae | 0.087 |  |
| Myoviridae | -0.137 | -0.171 |
| Siphoviridae | -0.137 | -0.171 |
| Anelloviridae | -0.361 |  |
| Betaflexiviridae | -0.361 | -0.474 |
| Bromoviridae | -0.361 | -0.474 |
| Closteroviridae | -0.361 | -0.474 |
| Paramyxoviridae | -0.361 | 0.132 |
| Picornaviridae | -0.361 | 3.466 |
| Adenoviridae | -0.585 | -0.777 |
| Alloherpesviridae | -0.585 | -0.171 |
| Baculoviridae | -0.585 | -0.171 |
| Bornaviridae | -0.585 |  |
| Endornaviridae | -0.585 |  |
| Filoviridae | -0.585 | -0.474 |
| Hantaviridae | -0.585 |  |
| Marnaviridae | -0.585 |  |
| Nairoviridae | -0.585 |  |
| Nimaviridae | -0.585 |  |
| Nodaviridae | -0.585 |  |
| Nudiviridae | -0.585 |  |
| Parvoviridae | -0.585 | 0.132 |
| Phenuiviridae | -0.585 |  |
| Pneumoviridae | -0.585 |  |
| Secoviridae | -0.585 | -0.777 |
| Tombusviridae | -0.585 | -0.777 |
| Unassigned | -0.585 |  |
| unclassified Arteriviridae | -0.585 |  |
| Alphaflexiviridae |  | 0.435 |
| Arenaviridae |  | -0.777 |
| Arteriviridae |  | -0.777 |
| Hepadnaviridae |  | -0.777 |
| Iflaviridae |  | -0.777 |
| Leviviridae |  | -0.777 |
| Luteoviridae |  | -0.777 |
| Microviridae |  | -0.777 |
| Peribunyaviridae |  | -0.777 |
| Polyomaviridae |  | -0.777 |
| Tymoviridae |  | -0.474 |
| Virgaviridae |  | 0.132 |

**Supplementary Table 3:**
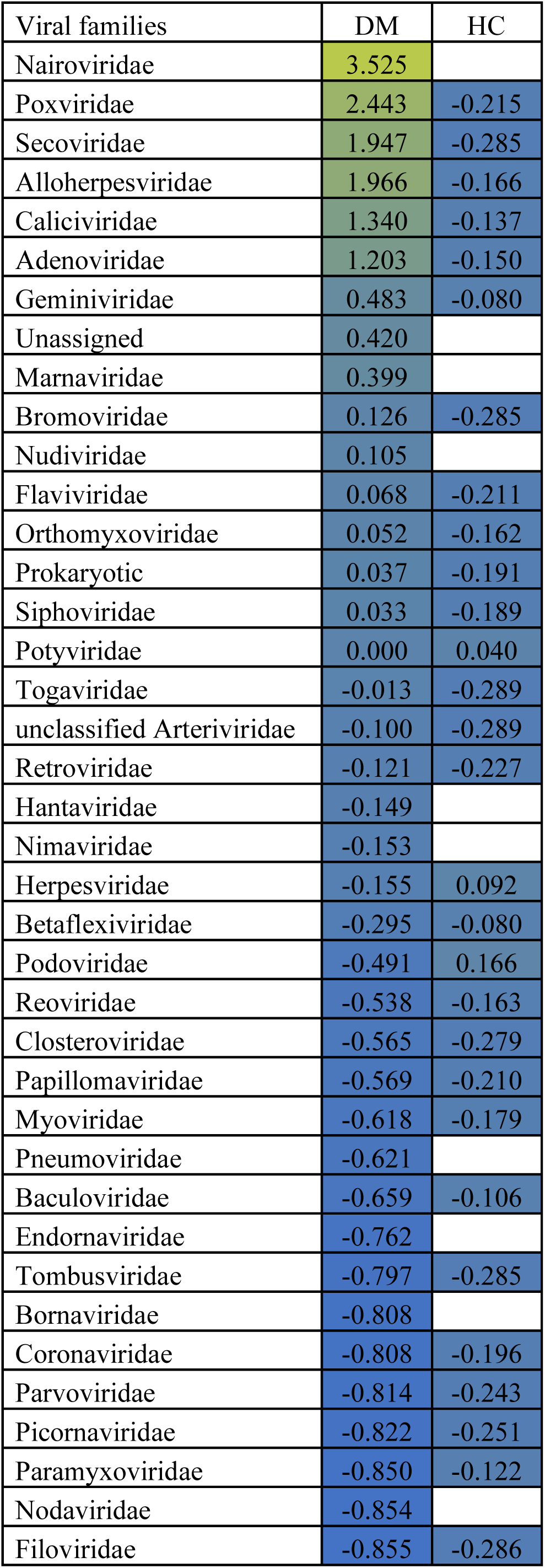
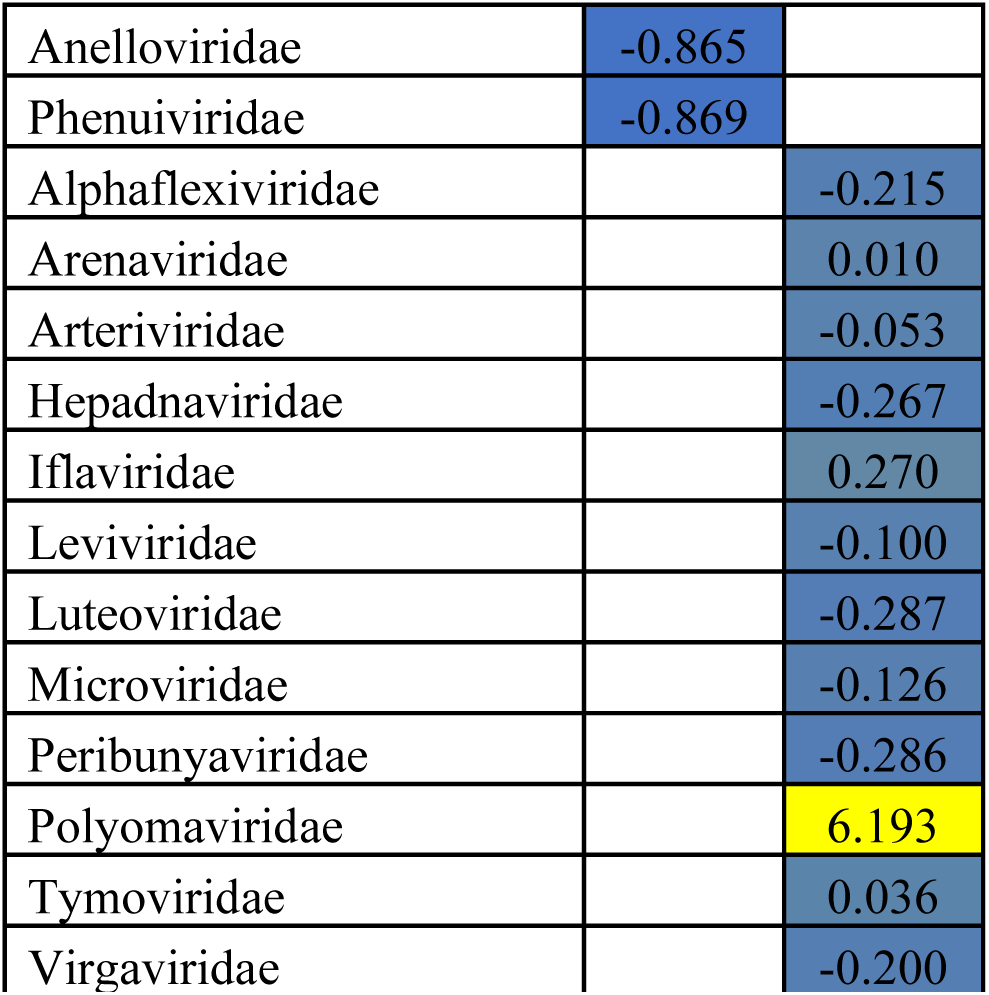
Mean NGSRe-norm of viral families identified in Dermatomyositis and healthy controls. The identified viral families are ordered based on decreasing mean NGSRe-norm in DM. Colour gradient is analogous to log10 transformed values of mean NGSRe-norm (Yellow: high, Blue: low). Missing values; Non-detected viral families.

| Viral families | DM | HC |
| --- | --- | --- |
| Nairoviridae | 3.525 |  |
| Poxviridae | 2.443 | -0.215 |
| Secoviridae | 1.947 | -0.285 |
| Alloherpesviridae | 1.966 | -0.166 |
| Caliciviridae | 1.340 | -0.137 |
| Adenoviridae | 1.203 | -0.150 |
| Geminiviridae | 0.483 | -0.080 |
| Unassigned | 0.420 |  |
| Marnaviridae | 0.399 |  |
| Bromoviridae | 0.126 | -0.285 |
| Nudiviridae | 0.105 |  |
| Flaviviridae | 0.068 | -0.211 |
| Orthomyxoviridae | 0.052 | -0.162 |
| Prokaryotic | 0.037 | -0.191 |
| Siphoviridae | 0.033 | -0.189 |
| Potyviridae | 0.000 | 0.040 |
| Togaviridae | -0.013 | -0.289 |
| unclassified Arteriviridae | -0.100 | -0.289 |
| Retroviridae | -0.121 | -0.227 |
| Hantaviridae | -0.149 |  |
| Nimaviridae | -0.153 |  |
| Herpesviridae | -0.155 | 0.092 |
| Betaflexiviridae | -0.295 | -0.080 |
| Podoviridae | -0.491 | 0.166 |
| Reoviridae | -0.538 | -0.163 |
| Closteroviridae | -0.565 | -0.279 |
| Papillomaviridae | -0.569 | -0.210 |
| Myoviridae | -0.618 | -0.179 |
| Pneumoviridae | -0.621 |  |
| Baculoviridae | -0.659 | -0.106 |
| Endornaviridae | -0.762 |  |
| Tombusviridae | -0.797 | -0.285 |
| Bornaviridae | -0.808 |  |
| Coronaviridae | -0.808 | -0.196 |
| Parvoviridae | -0.814 | -0.243 |
| Picornaviridae | -0.822 | -0.251 |
| Paramyxoviridae | -0.850 | -0.122 |
| Nodaviridae | -0.854 |  |
| Filoviridae | -0.855 | -0.286 |
| Anelloviridae | -0.865 |  |
| Phenuiviridae | -0.869 |  |
| Alphaflexiviridae |  | -0.215 |
| Arenaviridae |  | 0.010 |
| Arteriviridae |  | -0.053 |
| Hepadnaviridae |  | -0.267 |
| Iflaviridae |  | 0.270 |
| Leviviridae |  | -0.100 |
| Luteoviridae |  | -0.287 |
| Microviridae |  | -0.126 |
| Peribunyaviridae |  | -0.286 |
| Polyomaviridae |  | 6.193 |
| Tymoviridae |  | 0.036 |
| Virgaviridae |  | -0.200 |

